# Breast tumor microbiome regulates anti-tumor immunity and T cell-associated metabolites

**DOI:** 10.1101/2024.10.29.620864

**Authors:** Chin-Chih Liu, Dennis Grencewicz, Karthik Chakravarthy, Lin Li, Ruth Liepold, Matthew Wolf, Naseer Sangwan, Alice Tzeng, Rebecca Hoyd, Sachin R. Jhawar, Stephen R. Grobmyer, Zahraa Al-Hilli, Andrew P. Sciallis, Daniel Spakowicz, Ying Ni, Charis Eng

## Abstract

**Background:** Breast cancer, the most common cancer type among women, was recently found to contain a specific tumor microbiome, but its impact on host biology remains unclear. CD8^+^ tumor-infiltrating lymphocytes (TILs) are pivotal effectors of anti-tumor immunity that influence cancer prognosis and response to therapy. This study aims to elucidate interactions between CD8^+^ TILs and the breast tumor microbiome and metabolites, as well as how the breast tumor microbiome may affect the tumor metabolome.

**Methods:** We investigated the interplay among CD8^+^ TILs, the tumor microbiome, and the metabolome in a cohort of 46 breast cancer patients with mixed subtypes (Cohort A). We characterized the tumor metabolome by mass spectrometry and CD8^+^ TILs by immunohistochemistry. Microbiome composition and T cell gene transcript levels were obtained from data from our previous study, which utilized 16S rRNA gene sequencing and a targeted mRNA expression panel. To examine interactions between intratumoral *Staphylococcus* and specific breast cancer subtypes, we analyzed RNA sequencing data from an independent cohort of 370 breast cancer patients (Cohort B). We explored the functions of the tumor microbiome using mouse models of triple-negative breast cancer (TNBC).

**Results:** In tumors from Cohort A, the relative abundance of *Staphylococcus* positively correlated with the expression of T cell activation genes. The abundances of multiple metabolites exhibited significant correlations with CD8^+^ TILs, of which NADH, γ-glutamyltryptophan, and γ-glutamylglutamate displayed differential abundance in *Staphylococcus*-positive versus *Staphylococcus*-negative breast tumors. In a larger breast cancer cohort (Cohort B), we observed positive correlations between tumoral *Staphylococcus* and CD8^+^ TIL activity exclusively in TNBC. Preclinical experiments demonstrated that intratumoral administration of *S. aureus*, the predominant species of *Staphylococcus* in human breast tumors, resulted in a depletion of total NAD metabolites, and reduced the growth of TNBC tumors by activating CD8^+^ TILs.

**Conclusions:** We identified specific metabolites and microbial taxa associated with CD8^+^ TILs, delineated interactions between the breast tumor microbiome and metabolome, and demonstrated that intratumoral *Staphylococcus* influences anti-tumor immunity and TIL-associated metabolites. These findings highlight the role of low-biomass microbes in tumor tissues and provide potential biomarkers and therapeutic agents for breast cancer immunotherapy that merit further investigation.

## BACKGROUND

Breast cancer (BC) is the most prevalent cancer globally among women.(1) Treatment and prognosis hinge substantially on the cancer subtype, which is defined by the presence of the surface proteins estrogen receptor (ER), progesterone receptor (PR), and human epidermal growth factor receptor 2 (HER2). These proteins play pivotal roles in cancer initiation and progression and represent primary targets for therapeutic interventions. Triple-negative breast cancer (TNBC), characterized by the absence of these markers, is the most aggressive BC subtype. It is predominantly treated with chemotherapy and has a high recurrence rate. In the past decade, immunotherapies, particularly immune checkpoint inhibitors (ICIs), have become a transformative cancer treatment. Among BC subtypes, TNBC has demonstrated the best response to immunotherapy due to its higher immunogenicity, leading to the FDA approval of ICIs for certain TNBC patients.(2, 3) However, the relatively low response rate and therapeutic resistance remain major challenges of ICI therapy for TNBC, necessitating further investigation into the regulation of anti-tumor immunity.

CD8^+^ tumor-infiltrating lymphocytes (TILs) are critical effectors of the anti-tumor immune response and immunotherapy targets. The enrichment and activation of CD8^+^ TILs in BC are associated with improved treatment outcomes and prognosis.(4–6) Furthermore, using CD8^+^ TILs in adoptive cell therapy has shown efficacy against various cancers. Consequently, understanding the regulatory mechanisms of CD8^+^ TILs could yield novel therapeutic strategies and biomarkers for BC treatment. TIL activities can be inhibited by molecular products derived from the altered tumor metabolism, such as lactate generated by enhanced aerobic glycolysis (known as the Warburg effect) and oncometabolites (including D-2-hydroxyglutarate, succinate, and fumarate) accumulated by TCA cycle dysregulation.(7) To comprehensively decipher regulation of TILs by tumoral metabolites, it requires systematic profiling of the tumor metabolome alongside TIL analysis.

The function of CD8^+^ TILs is also influenced by the microbial communities present within tumors, referred to as the tumor microbiome. Since the majority of microbes in the human body predominantly reside in the gastrointestinal tract, tumors originating from these tissues can exhibit relatively high microbial biomass.(8) This tumor microbiome can impact cancer phenotypes through the modulation of reactive oxygen species, DNA damage responses, signaling pathways, inflammation, and metabolism.(9–12) For instance, in colorectal cancer, increased levels of intratumoral *Fusobacterium nucleatum* impair TIL function and modulate innate immune cells, indicating substantial influences of the tumor microbiome on anti-tumor immunity.(13) Notably, recent studies have suggested that microbes are also present in various types of tissue and solid tumors previously considered sterile, including glioblastoma, ovarian, bone, and breast tumors, albeit at much lower abundance than gastrointestinal tumors.(14–16) In these tumors with low microbial biomass, the impact of the tumor microbiome remains poorly understood.

Given that the tumor microbiome has shown strong correlations with prognosis and responses to treatment, including immunotherapy in certain cancer types, elucidating the composition and functions of the breast tumor microbiome may provide valuable insights into novel therapeutic strategies and prognostic markers.(17–20) Tzeng et al. showed that breast tumors have lower microbial diversity and reduced abundance of specific bacterial taxa, including *Staphylococcus* and *Streptococcus*, compared to normal breast and tumor-adjacent tissues.(16) Preclinical studies suggested that bacteria within breast tumors, such as *Bacteroides fragilis and Fusobacterium nucleatum*, could contribute to breast cancer development and metastasis.(21–23) Further, clinical evidence indicates that antibiotic use correlates with increased breast cancer risk and worse prognosis, suggesting the potential presence of beneficial microbes that could antagonize the development and progression of breast cancer and/or improve treatment efficacy.(24–26) However, it remains unclear whether there are tumor-suppressive bacteria localized within breast tumors that could activate CD8^+^ TILs and influence immunomodulatory metabolites.

This study examines interactions among CD8^+^ TILs, the tumor microbiome, and the metabolome in human breast tumors within a BC cohort without subtype stratification (Cohort A). We then explore the potential functions of *Staphylococcus* in tumors of specific BC subtypes in an independent, larger BC cohort (Cohort B). Beyond these correlative analyses, we employ murine models of TNBC, including 4T1 and EO771, to validate the functions of tumor-localized bacteria. Our results underscore the significant impact of the breast tumor microbiome on anti-tumor immune responses and CD8^+^ TIL-associated metabolites, which has implications for improving and designing cancer immunotherapy.

## METHODS

### Patient enrollment and tissue collection

Fresh-frozen breast tissues comprising Cohort A were acquired as described previously from three biorepositories, including the Cleveland Clinic Breast Center Microbiome Biorepository, Cooperative Human Tissue Network, and Case Comprehensive Cancer Center Human Tissue Procurement Facility.(16) Specimens were obtained using standard biorepository protocols from female patients who provided written informed consent. Breast tumor samples were obtained from oncologic surgery performed at Cleveland Clinic (n=16), University Hospitals Cleveland Medical Center (n=9), and The Ohio State University (n=21). Non-malignant breast tissues were collected from reduction mammoplasty performed at University Hospitals Cleveland Medical Center (n=10), Hospital of the University of Pennsylvania (n=7), University of Virginia Medical Center (n=7), and The Ohio State University (n=1). A pathologist verified that tissue samples from patients without breast cancer (healthy controls) to be free of malignant cells. Histopathological data were compiled from pathology reports. Breast cancer staging was standardized using the American Joint Committee on Cancer/Union for International Cancer Control 8th edition TNM pathologic stage criteria. Specimens were flash-frozen and stored at −80 °C until further processing. This study was approved by the Cleveland Clinic institutional review board (IRB #14-774 and 17-791). For Cohort B, 370 BC patients enrolled in a prospective cohort study (IRB protocol 2013H0199) at Ohio State University. Surgical specimens or tissue biopsies were sterilely collected, frozen in liquid nitrogen, and stored at -80 °C until further processing. Processed clinical and expression data were shared with researchers under an Ohio State University IRB-approved honest broker protocol (2015H0185).

### RNAseq data generation and processing

Fresh-frozen tissue was subjected to research use only (RUO) grade RNA sequencing at HudsonAlpha (Huntsville, AL) or Fulgent Genetics (Temple City, CA). Qiagen RNAeasy plus mini kit was used to isolate nucleic acid and fragmented to an average insert size of 216 bp.

Exons were enriched using the Illumina TruSeq RNA Exome with single library hybridization, followed by cDNA synthesis, library preparation, and sequencing to a depth of 50M paired-end reads. Adapters were trimmed via k-mer matching, followed by quality trimming and filtering, contaminant filtering, sequence masking, GC filtering, length filtering, and entropy filtering. Cleaned reads were processed for gene expression, microbe identification, and contaminant filtering using exotic v2.1.(27) Gene expression signatures were calculated using tnesig.(28) Immune cell abundances were estimated with CIBERSORTx.(29)

### RNAscope-fluorescence *in situ* hybridization (FISH)

All RNAscope-FISH was manually performed with the RNAscope Multiplex Fluorescent Reagent Kit v2 from Advanced Cell Diagnostics (ACD, Newark, CA). In brief, slides of human breast tumor tissues were baked in a 60°C oven for 2 hours and then deparaffinized in accordance with ACD recommendations. Target retrieval was performed with RNAscope Target Retrieval Reagents (PN 322000; ACD) warmed to 99°C in an Oster steamer for 15 minutes. Tissue was then subject to protease digestion with Protease Plus (PN 322331; ACD) and placed in a HybEZ II oven (PN 321710; ACD) for 30 minutes at 40°C. The protease was rinsed off with DI water. EB-16s-rRNA-C2 probe (464461-C2; ACD) was applied to the tissue and hybridized in a HybEZII oven for 2 hours at 40°C. Slides were then rinsed with RNAscope Wash Buffer (PN 310091; ACD). Amps 1-3 and Channel 1 from the Multiplex Fluorescent Detection Kit v2 (PN 323110; ACD) were applied to tissue in sequence and incubated in a HybEZII oven at 40°C for the duration recommended by ACD. The probe was visualized with Opal 570 (FP1488001KT; Akoya Biosciences) diluted in RNAscope Multiplex TSA Buffer (322809; ACD). Tissue was then stained with RNAscope Multiplex FL v2 DAPI (323108; ACD) and coverslipped using ProLong Gold antifade aqueous mounting media (P36930; Invitrogen). Images were acquired at 40X magnification on the Vectra Polaris Quantitative Pathology Imaging System.

### Immunohistochemistry and image analysis

The formalin-fixed, paraffin-embedded human breast tumor tissues were sectioned at 5 μm. The CD8^+^ and FoxP3^+^ cells present in these human tissues were identified by immunohistochemistry double stain using a DISCOVERY ULTRA automated stainer (Roche) followed by analysis using CaloPix, a computational pathology diagnostic software powered by artificial intelligence (Tribun Health, Paris). Details of these procedures were described in our previous study with a modification where CD8 was visualized by a DISCOVERY Yellow detection kit (Roche #760-228) after being stained with the primary and secondary antibodies.(16) CaloPix after trained by our images was used to identify CD8^+^ and FOXP3^+^ cells within tumor tissues. Identification of tumor-infiltrating CD8^+^ and FOXP3^+^ cells by CaloPix was validated by a breast pathologist.

Mouse tumor samples were prepared and subjected to immunohistochemistry based on the procedure described in our previous study with modifications as follows.(30) The primary antibody used for CD8 (Cell Signaling, #98941, 1:50) was diluted in a blocking solution and incubated at 4 °C overnight. The primary antibody for granzyme B (Abcam, #ab4059, 1:1500) was diluted in a blocking solution and incubated at room temperature for 1 h. After being washed with PBS (Dulbecco’s phosphate-buffered saline), tumor sections were incubated with horseradish peroxidase (HRP)-conjugated secondary antibody at room temperature for 30 min. The sections were subsequently stained with 3,3’-Diaminobenzidine (DAB) Substrate Kit and hematoxylin. Images were captured at 40X magnification on a Leica Aperio AT2 Slide Scanner and analyzed with QuPath digital pathology software. For the enumeration of stained cells, the function “Detect Positive Staining” was used to determine the positively stained pixels with DAB optical density (OD) value higher than 0.2 and 0.4 in the staining of granzyme B and CD8, respectively.

### Measurement of global untargeted metabolites

Untargeted metabolomics measurements were conducted at Metabolon (Morrisville, NC, USA) using Ultrahigh Performance Liquid Chromatography-Tandem Mass Spectroscopy (UPLC-MS/MS). Proteins were precipitated and removed from samples with methanol under vigorous shaking. The resulting extract was divided into five aliquots, dried, and reconstituted in solvents compatible with each of the methods containing a series of standards at fixed concentrations to ensure injection and chromatographic consistency. Among the five aliquots, two were analyzed by two separate reverse phase (RP)/UPLC-MS/MS methods with positive ion mode electrospray ionization (ESI), of which one was chromatographically optimized for more hydrophilic compounds while the other one was chromatographically optimized for more hydrophobic compounds. Two aliquots were analyzed by RP/UPLC-MS/MS and hydrophilic interaction liquid chromatography (HILIC)/UPLC-MS/MS, respectively, with negative ion mode ESI. One aliquot was reserved for backup. This strategy ensured maximal recovery and coverage of metabolites. All methods utilized a Waters ACQUITY ultra-performance liquid chromatography (UPLC) and a Thermo Scientific Q-Exactive high-resolution/accurate mass spectrometer interfaced with a heated electrospray ionization (HESI-II) source and Orbitrap mass analyzer operated at 35,000 mass resolution. Raw data was extracted, peak-identified and QC processed using Metabolon’s hardware and software. Compounds were identified by comparison to the Metabolon-maintained library based on the retention index, accurate mass match to the library +/-10 ppm, and the MS/MS score based on a comparison of the ions present in the experimental spectrum to the ions present in the library spectrum. The following QC and curation processes ensure accurate and consistent identification of true chemical entities and remove those representing system artifacts, misassignments, and background noise.

The raw values of the abundance of each metabolite were acquired according to the integrated area under the curve, which were then divided by the median of those samples to give each metabolite a median of one. The minimum value of each metabolite in the median scaled data is imputed for the missing values. Then the data were divided by the mass of input tissues and re-scaled to have a median equal to one. Finally, the metabolome data were transformed using the natural log function, which was used for statistical analysis.

### Canonical correlation analysis

We employed a sparse canonical correlation analysis (sCCA) to identify the potential crosstalk between tumoral microbiome and metabolites.(31) Canonical correlation analysis (CCA) has previously been suggested as a potential approach for performing integration analysis.(32) CCA is a technique used to identify weighted linear combinations of features from two distinct data modalities that exhibit significant association. This method aims to uncover relationships between variables measured on the same subjects. The typical CCA model provides non-zero weights to all features, potentially leading to overfitting when dealing with high-dimensional data. The problem of overfitting can be mitigated by incorporating a sparsity penalty into the CCA model, so enabling the inclusion of feature selection.(32)

The sCCA model can be mathematically represented as uTXTZv, subject to the following constraints: u 2≤1, v 2≤1, P1(u)≤c1, P2(v)≤c2, and uj=0 for all j∉Q1, and vj. The paired multi-omics datasets are represented by X and Z. The canonical vectors, u and v, include the weights assigned to each feature. The weighted linear combinations of features within each subject, Xu and Zv, are considered as the canonical scores. The P1 and P2 functions reflect the lasso penalty functions applied to the canonical variates. As a result, the values of u and v become sparse when the values of c1 and c2 are sufficiently tiny. The subsets of features in X and Z that exhibit significant univariate correlation with the phenotype are represented by Q1 and Q2. Conversely, features that do not strongly associate with the phenotype are assigned weights of zero automatically. The model’s optimal tuning parameters were determined using 25 times of permutation.

We conducted sCCA using the PMA package in R to select a parsimonious linear combination of variables that maximizes the correlation between two multivariate datasets.(31) The first dataset comprised normalized and scaled tumoral microbiome, and the second comprised metabolomics measurements from matched samples. Regularization parameters for the sCCA analysis were chosen by a grid search consisting of 100 possibilities of L1 values ranging from 0 (representing the highest sparsity) to 1 (representing the lowest sparsity), with increments of 0.1. We chose the set of L1 values that resulted in the highest canonical correlation of the first variate for further examination.

### Cell culture

The sources, culture media, sterility, and authentication of 4T1 and EO771 cell lines were described in our previous study.(30)

### Bacterial culture

To acquire live suspensions of bacteria, *S. aureus* (BEI Resources Repository, # HM162), *S. epidermidis* (BEI Resources Repository, # HM118), *S. hominis* (ATCC, # 27844), *S. oralis* (ATCC, # 700233), *S. mitis* (BEI Resources Repository, # HM262), *C. acnes* (ATCC, # 6919), *P. aeruginosa* (ATCC, # 15692), and *B. Longum* (ATCC, # 15707) were inoculated into BHI media (Sigma, # 110493) supplemented with menadione (1 mg/l, MP Biomedicals, # 02102259), hematin (1.2 mg/l, Santa Cruz, # sc-207729), histidine (0.2 mM, Tokyo Chemical Industry, # H0149), and L-cysteine hydrochloride (0.5 g/l, J.T.Baker, # 2071-05) and incubated at 37 °C in an anaerobic chamber, except for *P. aeruginosa*, which is incubated in an aerobic shaker. *L. lactis* (ATCC, # 19435) was grown aerobically at 37 °C in the aforementioned media supplemented with 5% serum. *C. tuberculostearicum* (ATCC, # 35692) was grown aerobically at 37 °C in PYG media supplemented with menadione (0.315 mg/l), hematin (5 mg/l), histidine (0.834 mM), and L-cysteine hydrochloride (0.5 g/l). On the day of intratumoral injection, bacteria were pelleted down by centrifugation, washed, and resuspended in PBS at a concentration of 5 × 10^9^ CFU (colony-forming units) per ml, which was confirmed by enumeration of plated colonies.

### Tumor growth and treatment

4T1 and EO771 cells were trypsinized and resuspended in PBS without calcium and magnesium. 1 x 10^4^ of 4T1 cells and 2.5 x 10^5^ to 1 x 10^6^ of EO771 cells were injected into the fourth mammary fat pad of six to eight week old female BALB/c and C57BL/6J mice (The Jackson Laboratory), respectively. The number of injected cells was the same between different treatment groups of every experiment. After palpable tumors formed, mice were randomized with matching tumor volumes into case and control groups before treatment. Tumor volumes were measured by digital calipers and calculated using the ellipsoid formula (L x W x W/2). The day of the first bacterial injection was defined as day zero hereafter. To investigate the effects of intratumoral bacteria on the growth, tumor microenvironment, and total NAD level of EO771 tumors, 2 x 10^7^ CFU of bacteria were injected into palpable tumors on day zero and the tumors were isolated on day 10 for downstream analyses. To characterize the effects of intratumoral bacteria on 4T1 tumor growth, 1 x 10^8^ CFU of bacteria were injected into 4T1 tumors on days zero and day four, except for *P. aeruginosa*, which was injected at a dosage of 1 x 10^6^ CFU due to its systemic toxicity. To determine the influence of intratumoral bacteria on TILs in 4T1 tumors, 1 x 10^6^ CFU of bacteria were injected into tumors on day zero, and tumors were isolated for immune profiling on day 12. To test whether lymphocytes are required for the *S. aureus*-mediated inhibition of tumor growth, tumor-bearing mice were subjected to intraperitoneal injections of anti-CD4 (BioXCell, clone GK1.5, 150 μg), anti-CD8 (BioXCell, clone YTS 169.4, 150 μg), or isotype IgG control (BioXCell, clone LTF-2, 150 μg) one day before the intratumoral injection of 2 X 10^7^ CFU of *S. aureus*, which was followed by an additional two injections of the same neutralizing antibodies on days three and six.

### Immune cell dissociation and flow cytometry

Tumors were digested as described previously.(30) For the intracellular analysis of granzyme B, dissociated cells were additionally cultured for 4 hours with 50 nM phorbol 12-myristate 13-acetate (PMA, Sigma, # P1585), 500 nM ionomycin (Sigma, # I0634), and Brefeldin A (Biolegend, # 420601) before staining. Collected cells were stained with viability dye (Biotium, # 32018, 1:1000) for 30 minutes, blocked with anti-mouse CD16/32 antibody (Biolegend, # 156604, 1:100) in staining buffer (PBS with 0.5% Bovine Serum Albumin (BSA), 2mM EDTA, and 0.05% NaN_3_) for 20 minutes, and then stained with antibodies targeting cell surface markers for 30 minutes on ice. After two washes with staining buffer, cells were resuspended in fixation buffer at room temperature for 60 min and permeabilized using the permeabilization buffer according to the manufacturer’s instructions (Biolegend, # 424401; or Thermo Fisher Scientific, # 88-8824-00 for granzyme B analysis). Cells were then stained for 45 min at room temperature with antibodies targeting intracellular markers. Samples were analyzed using a SONY ID7000™ spectral cell analyzer. The resulting FCS data files were then analyzed by FlowJo™ Software v10.8.0 (BD Life Sciences) with the gating strategy shown in Supplementary Fig. 10. The following antibodies were used in this study: CD45 (BD Biosciences, clone 30-F11), CD11b (Thermo Fisher Scientific or BD Biosciences, clone M1/70), Ly-6C (Biolegend, clone HK1.4), Ly-6G (BD Biosciences, clone 1A8), I-A/I-E (BD Biosciences, clone M5/114.15.2), CD3 (Biolegend, clone 17A2), CD11c (BD Biosciences, clone N418), CD206 (Biolegend, clone C068C2), CD4 (BD Biosciences, clone GK1.5), F4/80 (Biolegend, clone BM8), CD8a (Biolegend, clone 53-6.7), phosphorylated STAT3 (Thermo Fisher Scientific, clone LUVNKLA), Granzyme B (Thermo Fisher Scientific, clone NGZB).

### NAD measurement

10 days after bacterial injection, EO771 tumors were isolated, washed with ice-cold PBS, and cut into small pieces using sterile dissection tools inside a positive flow cell culture hood. The total level of NAD^+^ and NADH was measured by the NAD/NADH Quantification Kit (Sigma, # MAK037). In brief, 15 to 30 mg of tumor pieces were homogenized with 700 μl of NADH/NAD Extraction Buffer in a Dounce homogenizer on ice. Following vigorous vortexing and centrifugation, the supernatant was filtered through a 10 kDa cut-off spin filter (Abcam # ab93349) to remove potential NAD^+^/NADH-consuming enzymes. After the addition of NAD Cycling Enzyme and NADH Developer, the absorbance at 450 nm was measured using a microplate reader (BioTek Synergy H1). The protein concentration of the samples before filtration was determined by a bicinchoninic acid (BCA) protein assay (ThermoFisher, # 23227) for the normalization of the results.

### Data normalization and statistical analyses

The data on microbiome composition and expression levels of 579 immunity-related genes in the human breast tumors comprising Cohort A were retrieved from the data of our previous study.(30) Only the bacterial genera that were identified in more than 5% of breast tissue samples were included for downstream analysis. The raw values of the abundance of each bacterial genera were normalized to the mass of input tissues used for 16S rRNA sequencing. The further normalization was performed based on the formula log10(RCn X ∑xN + 1), where RC represents the number of counts i for a sample, n is the number of sequences in a sample, the sum of x is the total number of counts in the table, and N is the total number of samples.(33)

Statistical analyses were performed with R version 4.3.1 and Prism version 10.1.1 (GraphPad). Unpaired two-tailed Student’s t-test or Mann-Whitney u test was used to compare experiments with two groups. One-way analysis of variance (ANOVA) was used to compare experiments with greater than two groups. To compare tumor growth, Two-way ANOVA with Sidak’s and Tukey’s multiple comparison tests were performed with two and greater than two groups, respectively. *P* < 0.05 was used as a threshold of statistical significance. Spearman’s correlation was applied to analyze the associations between microbiome and TILs, microbiome and T cell genes, metabolites and TILs, and metabolites and T cell genes. Sparse canonical correlation analysis was used to identify the tumoral metabolites associated with specific tumor microbes. A p-value without adjustment for multiple comparisons was used to determine the significance of statistical analyses in identifying hot and cold metabolites to minimize type II errors. To reduce type I errors, two statistical approaches, including Spearman’s correlation and Mann-Whitney U test, were simultaneously applied, and only the metabolites identified with significant unadjusted p-values under both statistical analyses were defined as hot or cold metabolites. Refer to Results for details.

## RESULTS

### Patient characteristics of Cohort A

To examine interrelationships among CD8^+^ TILs, the tumor microbiome, and metabolome, we collected multidimensional data derived from 46 human breast tumors from 46 breast cancer patients encompassing different BC subtypes (Cohort A). The microbiome composition of all 46 tumors from Cohort A has been profiled by 16S rRNA gene sequencing, and transcript levels of immunity-related genes in 41 tumors from this cohort have been analyzed by Nanostring in our earlier investigation.(16) Using additional tumor tissues from the same cohort, this study further conducted tumor metabolome profiling and immunohistochemistry using anti-CD8 and anti-FoxP3 antibodies. For metabolome analysis, an additional 25 non-malignant healthy breast tissues obtained from reduction mammoplasty were included as healthy controls. Detailed demographic and clinicopathologic characteristics of patients are shown in Table 1.

**Table 1.**
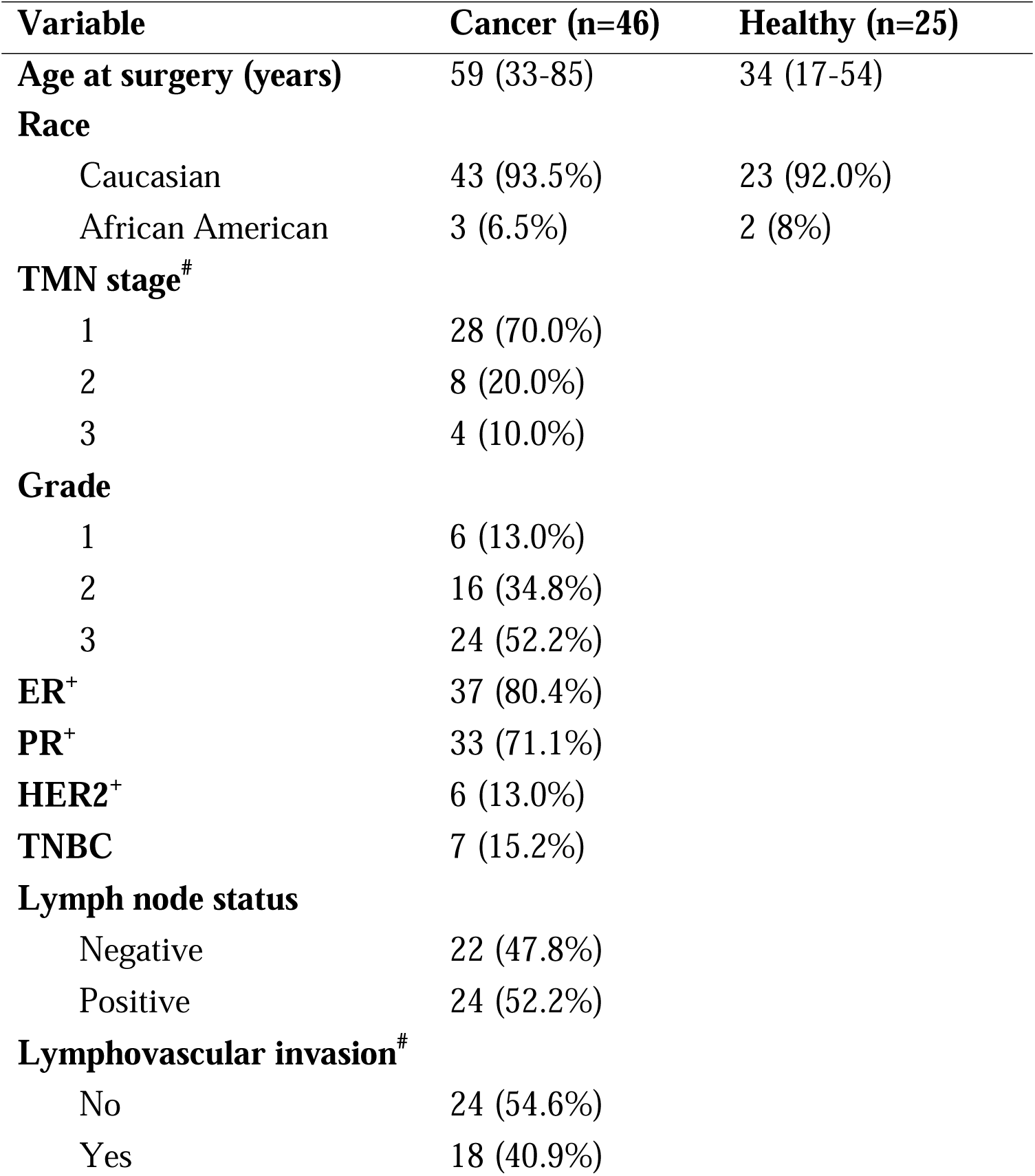
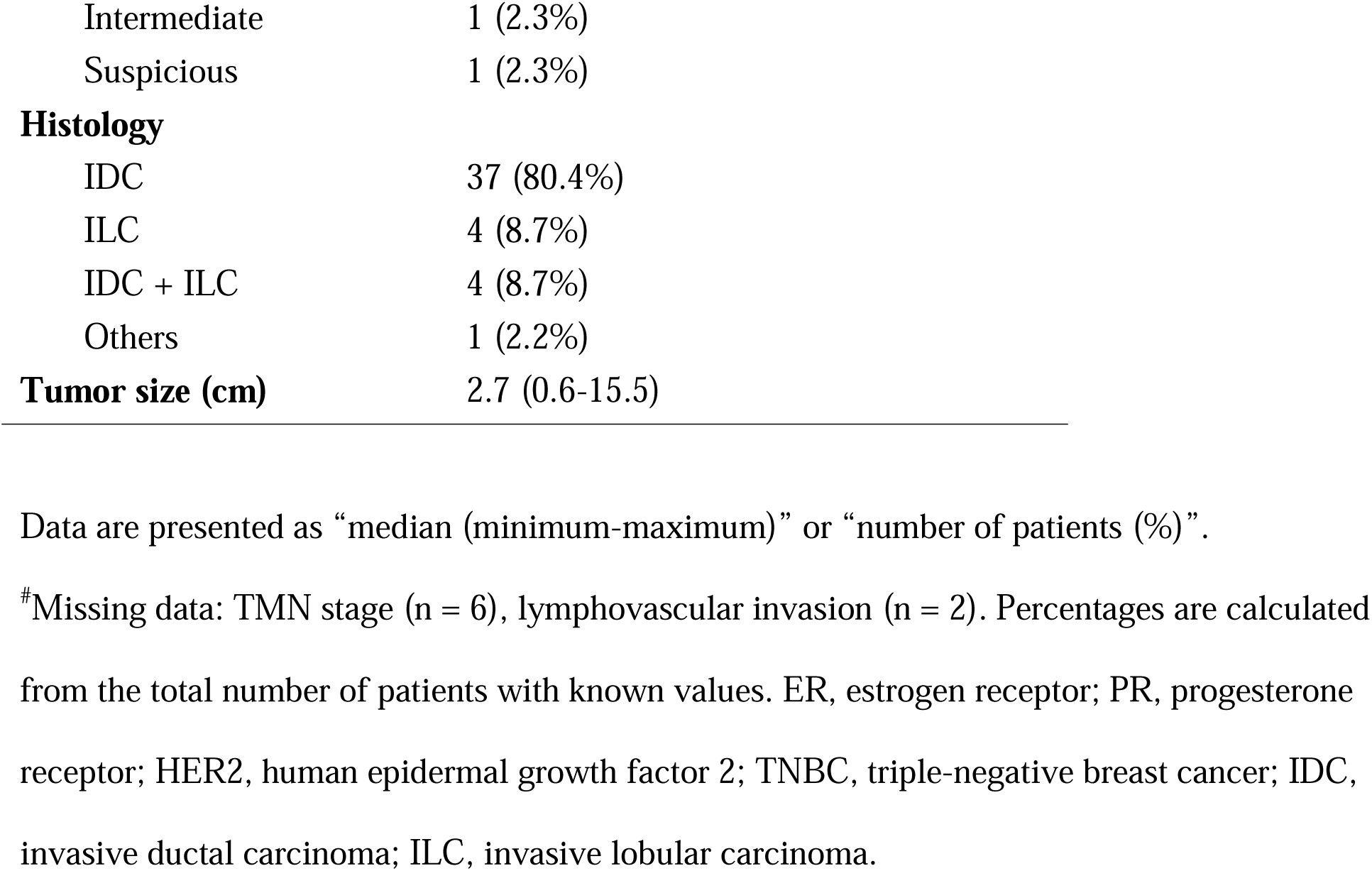
Patient characteristics (Cohort A)

| Variable | Cancer (n=46) | Healthy (n=25) |
| --- | --- | --- |
| Age at surgery (years) | 59 (33-85) | 34 (17-54) |
| **Race** |  |  |
| Caucasian | 43 (93.5%) | 23 (92.0%) |
| African American | 3 (6.5%) | 2 (8%) |
| **TMN stage<sup>#</sup>** |  |  |
| 1 | 28 (70.0%) |  |
| 2 | 8 (20.0%) |  |
| 3 | 4 (10.0%) |  |
| **Grade** |  |  |
| 1 | 6 (13.0%) |  |
| 2 | 16 (34.8%) |  |
| 3 | 24 (52.2%) |  |
| **ER<sup>+</sup>** | 37 (80.4%) |  |
| **PR<sup>+</sup>** | 33 (71.1%) |  |
| **HER2<sup>+</sup>** | 6 (13.0%) |  |
| **TNBC** | 7 (15.2%) |  |
| **Lymph node status** |  |  |
| Negative | 22 (47.8%) |  |
| Positive | 24 (52.2%) |  |
| **Lymphovascular invasion<sup>#</sup>** |  |  |
| No | 24 (54.6%) |  |
| Yes | 18 (40.9%) |  |

|  |  |
| --- | --- |
| Intermediate | 1 (2.3%) |
| Suspicious | 1 (2.3%) |
| **Histology** |  |
| IDC | 37 (80.4%) |
| ILC | 4 (8.7%) |
| IDC + ILC | 4 (8.7%) |
| Others | 1 (2.2%) |
| **Tumor size (cm)** | 2.7 (0.6-15.5) |

Data are presented as “median (minimum-maximum)” or “number of patients (%)”.
#Missing data: TMN stage (n = 6), lymphovascular invasion (n = 2). Percentages are calculated from the total number of patients with known values. ER, estrogen receptor; PR, progesterone receptor; HER2, human epidermal growth factor 2; TNBC, triple-negative breast cancer; IDC, invasive ductal carcinoma; ILC, invasive lobular carcinoma.

### *Staphylococcus* is associated with activation of TILs in human breast tumors

To verify the presence of bacteria in human breast tumors beyond relying on sequencing data alone, we performed RNAscope-FISH using a probe targeting the bacterial 16S rRNA gene, which displayed punctate bacteria scattered across tumor cells (Fig. 1a). To identify which bacterial taxa within the breast tumor microbiome are associated with TILs, we quantified the abundance of CD8^+^ and FoxP3^+^ TILs and retrieved expression levels of genes indicative of CD8^+^ T cell activation from our prior data, including GZMA/B/K and IFNG. Correlation analysis showed that among 18 predominant bacterial genera of the tumor microbiome, *Acinetobacter* and *Dietzia* were inversely correlated with the abundance of FoxP3^+^ cells, and *Pelomonas* was negatively associated with GZMA level (Fig. 1b, c). Notably, *Staphylococcus* was the only genus that exhibited statistically significant positive associations with transcript levels of GZMA, B, and K (Fig. 1c). Of all the bacteria genera, *Staphylococcus* also displayed the strongest correlation with IFNG level. To further categorize *Staphylococcus* at the species level, we reanalyzed the 16S rRNA gene sequencing data and discovered that *S. aureus* is the most abundant species of *Staphylococcus* identified in human breast tumors (supplementary Fig. 1). These results suggest a potential role of *Staphylococcus* and, in particular, *S. aureus* in stimulating CD8^+^ T cells locally within human breast tumors.

**Figure 1.**
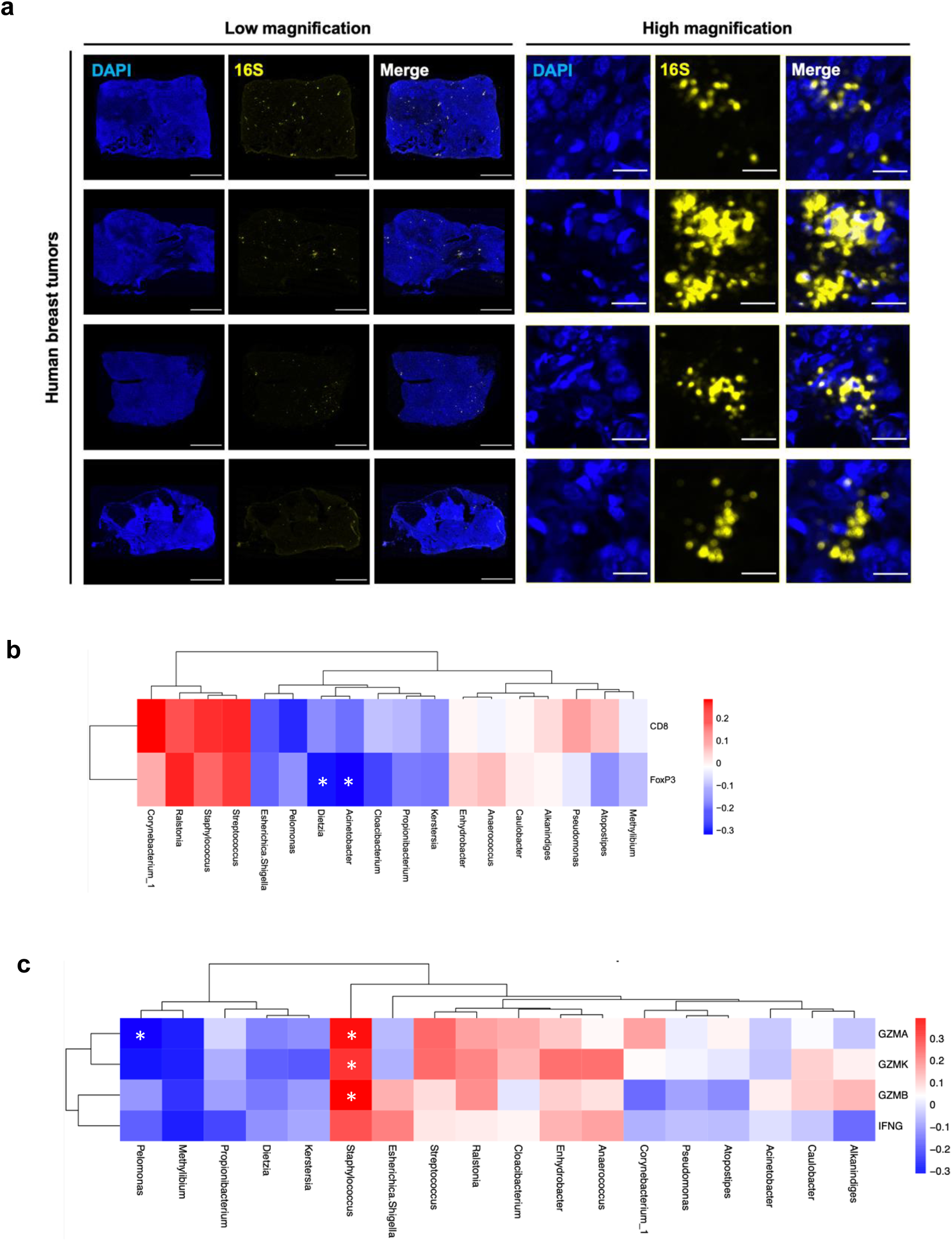
Presence of tumoral bacteria and their correlations with TIL abundance and activation genes. **a** RNAscope-fluorescence in situ hybridization (FISH) images of human breast tumors using a probe targeting the bacterial 16S gene (yellow). Nuclei were stained with DAPI (blue). Scale bars are shown in white at 2000 and 25 μm in the images of low and high magnification, respectively. **b, c** Heatmaps displaying correlations between the abundance of bacterial genera and CD8^+^ and FoxP3^+^ cells (**b**) as well as the genes indicative of T cell activation, including GZMA/B/K and IFNG (**c**). Spearman’s correlation (**b, c**). \**P* < 0.05.

### Identification of tumoral metabolites associated with CD8^+^ TILs

To investigate associations between tumoral metabolites and CD8^+^ TILs, we conducted global metabolome profiling of human breast tumors alongside healthy breast tissues as non-malignant controls. Using untargeted metabolomics, we identified over 900 diverse metabolites (supplementary Fig. 2a). As expected, breast tumors were metabolically distinct from healthy breast tissues and exhibited enhanced metabolic activity across various pathways (supplementary Fig. 2b, c, d). Tumor tissues displayed a higher ratio of glutamate to glutamine, suggesting a higher level of glutaminolysis (supplementary Fig. 2e).(34) The top 25 metabolites enriched in breast tumors and healthy breast tissues are listed in supplementary Table 1 and 2, respectively. Breast tumor metabolites correlated with key clinicopathologic features, including cancer stage, histological grade, tumor size, lymph node status, histological subtypes, and patient age (supplementary Fig. 3a-e). The glutamate-to-glutamine ratio was also different between the stages and subtypes of breast cancer (supplementary Fig. 4a, b). Based on intratumoral CD8^+^ cell density quantified using immunohistochemistry, we stratified breast tumors into hot and cold categories, corresponding respectively to tumors with relatively higher and lower densities of CD8^+^ T cells (Fig. 2a). PCA results showed subtle differences in the metabolome between hot and cold tumors (Fig. 2b). We then employed two statistical methods to identify metabolites associated with CD8^+^ TILs. The Mann–Whitney U test highlighted metabolites with differential abundance between hot and cold tumors, while Spearman’s rank correlation identified metabolites correlated with CD8^+^ cell density. A total of 39 metabolites exhibited significant p-values in both analyses (Fig. 2c), of which 29 and 10 metabolites were enriched in hot and cold tumors, respectively (henceforth referred to as hot and cold metabolites; Fig. 2d). Hot metabolites encompassed various lipids including medium/long chain fatty acids and N-acylethanolamines (NAEs), compounds of the γ-glutamyl cycle, metabolic aspirin derivatives, and microbiota-derived 3-formylindole and imidazole propionate. Cold metabolites included succinate, bilirubin (E,E), secondary bile acids, nicotinamide adenine dinucleotide (NAD), and steroids including dehydroepiandrosterone sulfate (DHEA-S), androstenediol (3beta,17beta) disulfate, and pregnenediol sulfate (C21H34O5S). Notably, NAD^+^ and NADH emerged as the most significant cold metabolites that exhibited significantly heightened abundance in cold tumors compared to hot tumors and healthy breast tissues (Fig. 2e, f). Additionally, several hot and cold metabolites including NAD^+^, NADH, taurolithocholate 3-sulfate, and 3-formylindole were found to associate with not only the abundance of CD8^+^ cells but also the expression levels of genes related to T cell activation, suggesting their associations with both CD8^+^ TIL abundance and activity (Fig. 2g).

**Figure 2.**
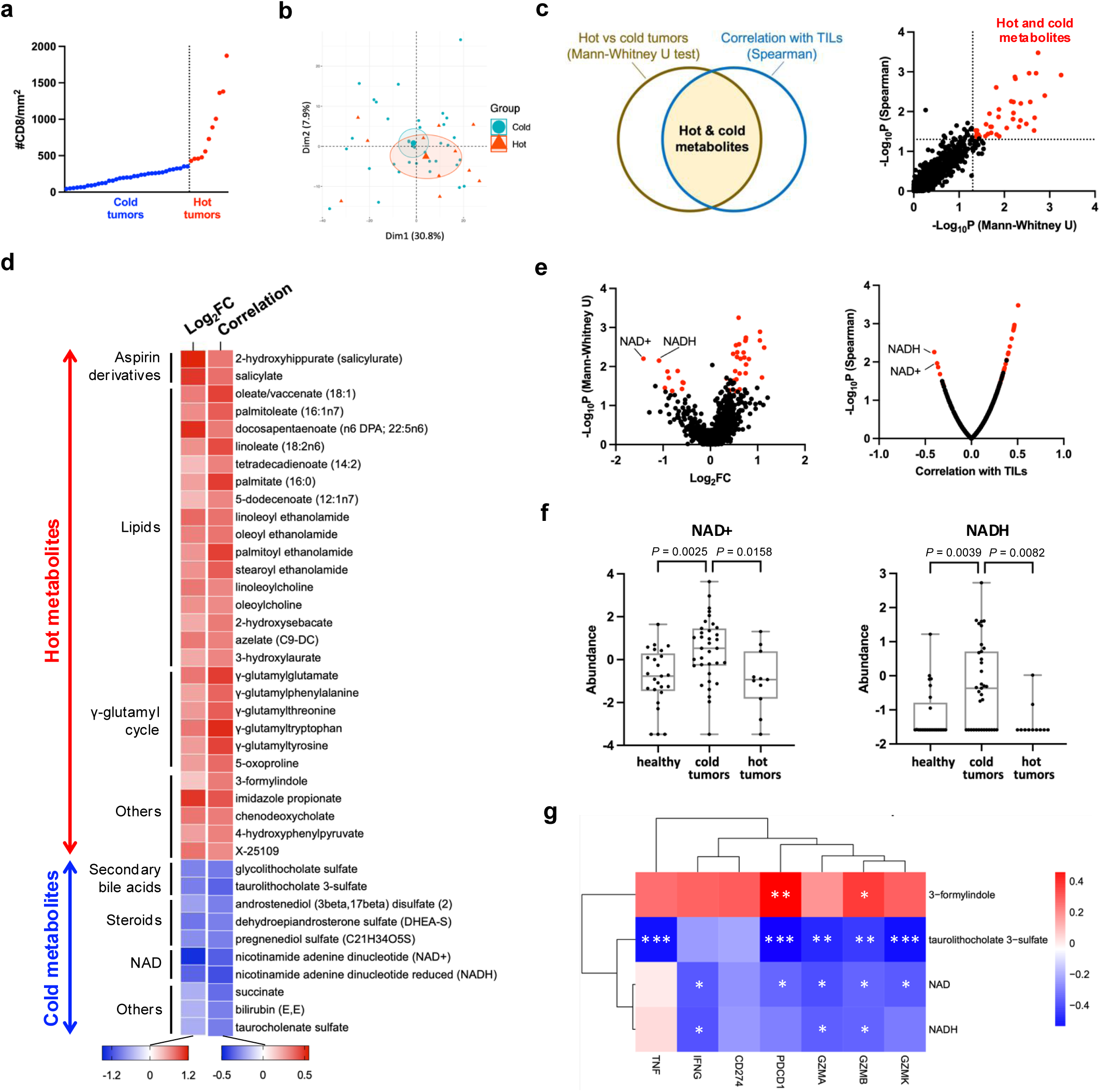
Identification of tumoral metabolites associated with CD8^+^ TILs. **a** Categorization of hot and cold tumors based on CD8^+^ cell density quantified using immunohistochemistry. **b** Principal component analysis (PCA) of metabolites in hot and cold tumors (n=11 and 35, respectively). **c** The left panel shows the two statistical approaches employed to identify hot and cold metabolites. The right figure depicts hot and cold metabolites that exhibit significant p-values in both analyses in red. **d** Heatmap showing the identified hot and cold metabolites. The color gradient represents the log_2_ fold change (FC) of each metabolite between hot and cold tumors (left column) and the correlation between each metabolite and CD8^+^ cell density (right column). **e** Volcano plot showing hot and cold metabolites (both highlighted in red) derived from Mann-Whitney U test (left) and Spearman’s rank correlation (right). **f** The abundance of NAD^+^ (left panel) and NADH (right panel) in healthy breast tissues (Healthy, n=25), cold tumors (n=35), and hot tumors (n=11). **g** Heatmap representing correlations between transcript levels of T cell-related genes and the abundance of NAD^+^, NADH, taurolithocholate 3-sulfate, and 3-formylindole. One-way analysis of variance (ANOVA) with multiple comparisons (**f**). Spearman’s correlation (**g**). \**P* < 0.05; \*\**P* < 0.01; \*\*\**P* < 0.001.

### *Staphylococcus* correlates with CD8^+^ TIL-associated tumoral metabolites

Our investigation then turned toward exploring the crosstalk between the tumoral microbiome and metabolites. Sparse canonical correlation analysis (sCCA) revealed that distinct bacterial genera associated with different patterns of metabolites (supplementary Fig. 5). *Anaerococcus* and *Methylibium* were the only genera significantly correlated with the ratio of glutamate to glutamine (supplementary Fig. 6). Since *Staphylococcus* is the only bacterial genus that exhibited significant correlation with T cell activity (Fig. 1c), we conducted a targeted analysis to determine whether *Staphylococcus* associates with the metabolites linked to CD8^+^ TILs and clinicopathological features of breast cancer. Our findings indicated that *Staphylococcus*-positive tumors displayed a decreased level of cold metabolite NADH, along with higher levels of hot metabolites γ-glutamyltryptophan and γ-glutamylglutamate (Fig. 3). Additionally, *Staphylococcus*-positive tumors demonstrated reduced levels of N-acetyl-aspartyl-glutamate (NAAG), dihydroorotate, and GDP as well as increased maleate, of which NAAG, dihydroorotate and maleate are compounds found to be associated with breast tumor, tumor size, and TNBC, respectively (supplementary Fig. 3, and supplementary Table 1). These results reveal specific associations between *Staphylococcus* and CD8^+^ TILs-associated metabolites in breast tumors.

**Figure 3.**
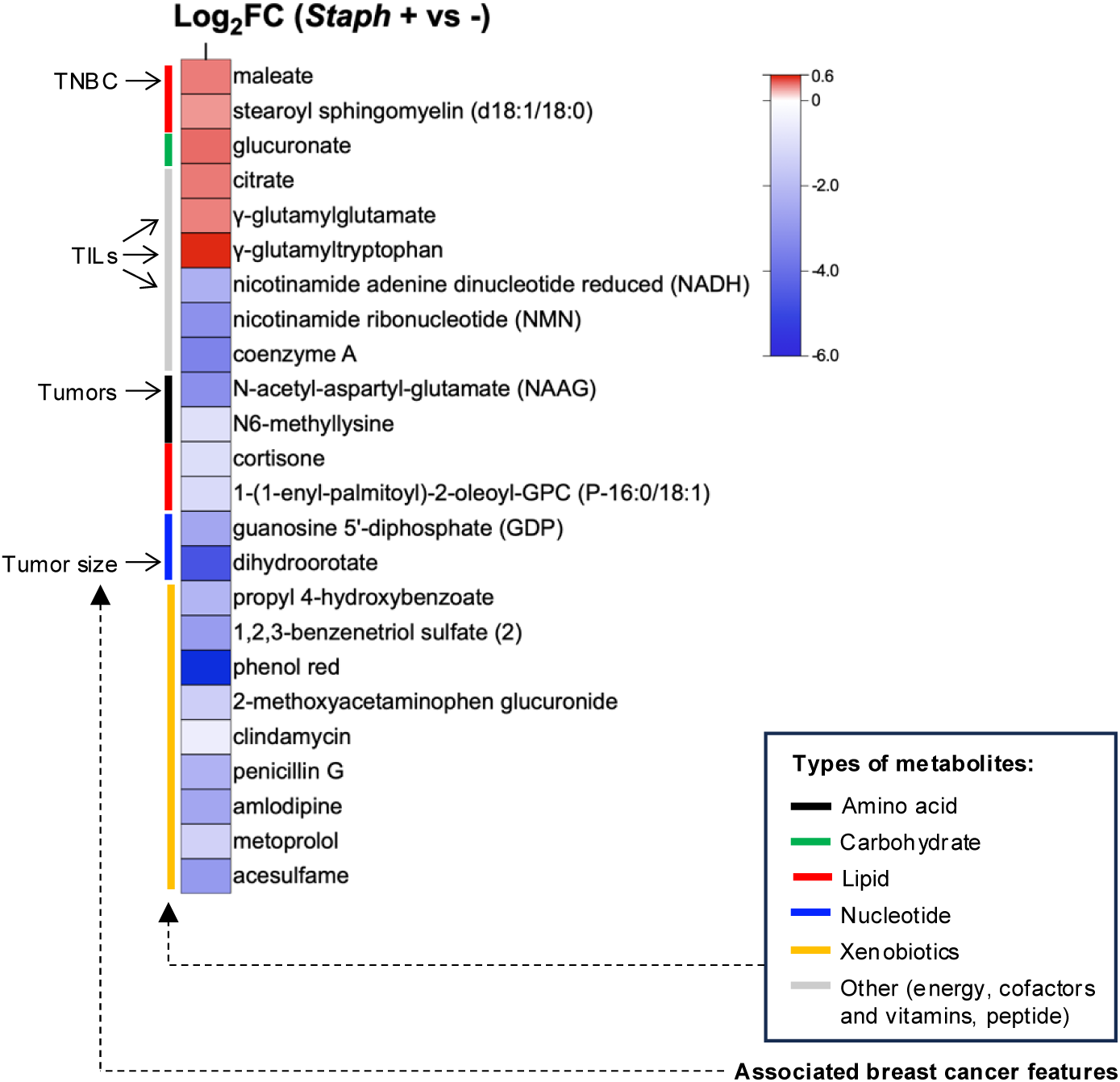
***Staphylococcus-*associated tumoral metabolites. a** Heatmap displaying differentially abundant metabolites in *Staphylococcus*-positive compared to *Staphylococcus*-negative (*Staph+* vs *-*) human breast tumors. The color gradient represents the log_2_ fold change (FC) relative to the *Staphylococcus*-negative group. Types of metabolites and their associated breast cancer features (detailed in Fig. 2, supplementary Fig. 3, and supplementary Table 1) are indicated on the left side of the heatmap. Unpaired two-tailed Student’s t-test.

### *Staphylococcus* is associated with CD8^+^ TIL activation in TNBC

To further elucidate the roles of *Staphylococcus* in different BC subtypes and prognosis, we analyzed the RNA sequencing data from an independent clinical cohort of breast tumor tissues collected from The Ohio State University Wexner Medical Center (Cohort B). This cohort comprised 370 breast cancer patients who did not receive neoadjuvant therapy before sample collection (Table 2). Among the various cancer subtypes, the proportion of *Staphylococcus*-positive tumors did not show significant differences (Fig. 4a). The percentage of Staphylococcus-positive tumors consistently exhibited a declining trend with advancing cancer stages across each cancer subtype, although this trend was not statistically significant (Fig. 4b). The presence of *Staphylococcus* in tumors was not significantly associated with prognosis in all BCs or in either subtype (TNBC or ER^+^/PR^+^ subtype), but it was associated with a slight, albeit non-significant, increase in the abundance of CD8^+^ TILs in TNBC (Fig. 4c, d). Importantly, *Staphylococcus*-positive TNBC exhibited a significant increase in CD8^+^ T cell activity compared to *Staphylococcus*-negative TNBC (Fig. 4e). Among the T cell activation-related genes, *Staphylococcus*-positive TNBC showed significant increases in levels of GZMB, PRF1, IL2RA (CD25), CD38, PDCD1, LCK, ZAP70, and TBX21 (T-BET), as well as non-significant trends toward elevated GZMA, GZMK, and IFNG (Fig. 4f). Intriguingly, *Staphylococcus*-positive

**Figure 4.**
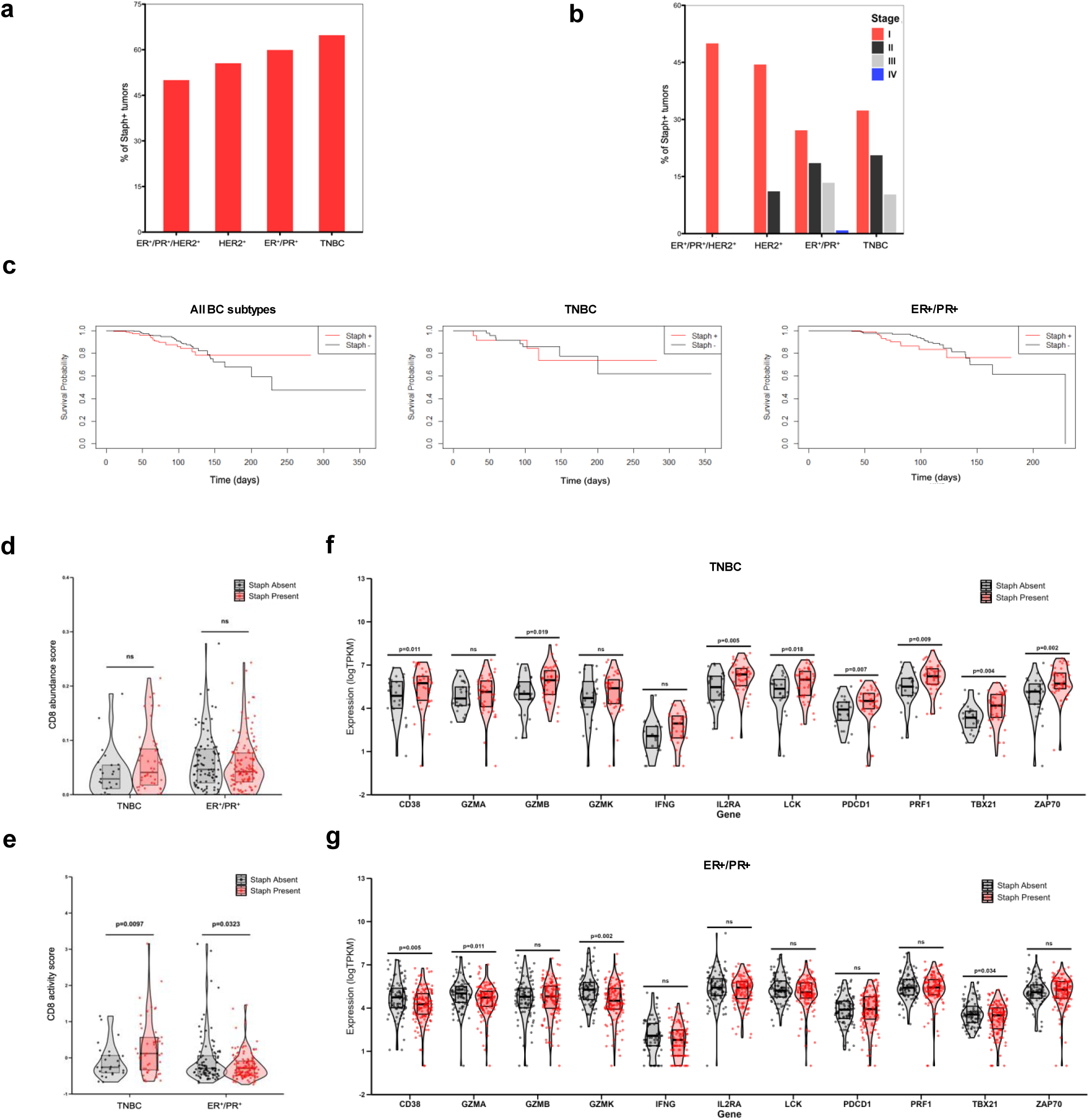
Associations between *Staphylococcus*, clinicopathological features, and CD8^+^ TILs. a, **b** Percentages of Cohort B human breast tumor samples identified as positive for *Staphylococcus* (Staph+ tumors) across cancer subtypes (**a**) and stages (**b**). **c** Kaplan–Meier survival curves illustrating overall survival among patients of all breast cancer subtypes (left panel), TNBC (middle panel), and ER+/PR+ subtype (right panel) stratified by the presence or absence of intratumoral *Staphylococcus* (Staph+ and Staph-, shown as red and black lines, respectively). All p-values > 0.05 by log-likelihood test. **d**, **e** Comparisons of CD8+ T cell abundance (**d**) and activity (**e**) in TNBC and ER+/PR+ tumors with and without the presence of *Staphylococcus* (shown as red and black dots, respectively). **f**, **g** Comparisons of T cells-related genes in TNBC (**f**) and ER+/PR+ tumors (**g**) with and without the presence of *Staphylococcus*. Z-score transformed signature scores, and log-transformed transcripts per kilobase-million (TPKM) counts were compared by t-test. ns, not significant.

**Table 2.**
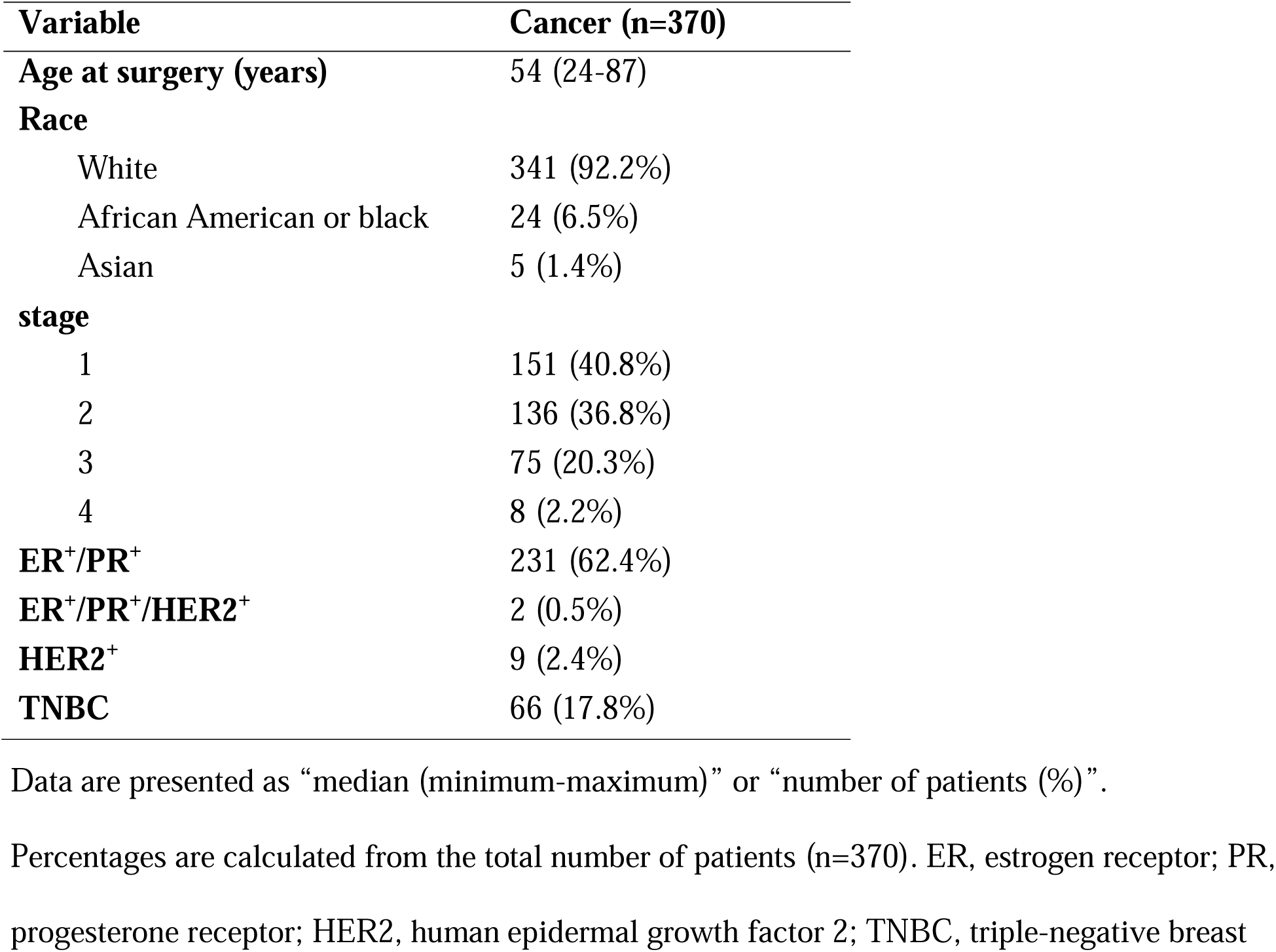
Patient characteristics (Cohort B)

ER^+^/PR^+^ tumors displayed reduced CD8^+^ T cell activity and significantly lower levels of GZMA, GZMK, CD38, and TBX21 compared to *Staphylococcus*-negative ER^+^/PR^+^ tumors (Fig. 4e, 4g). Furthermore, the presence of *Staphylococcus* was not associated with differences in host metabolic pathways related to NAD metabolism or the γ-glutamyl cycle in TNBC and ER^+^/PR^+^ subtypes, suggesting that *Staphylococcus* may directly regulate T cell-associated metabolites including NADH, γ-glutamyltryptophan, and γ-glutamylglutamate levels without altering associated host genes (Supplementary Fig. 7a, b). Collectively, these findings reveal differential interactions between tumor-localized *Staphylococcus* and CD8^+^ TIL activity in triple-negative and ER^+^/PR^+^ tumors, and indicated that *Staphylococcus* may activate CD8^+^ TILs in TNBC.

### Intratumoral *S. aureus* modulates CD8^+^ TIL-dependent anti-tumor immunity and total NAD level in TNBC models

To investigate whether *Staphylococcus* and other tumoral bacteria affect TIL activity and anti-tumor immunity, functional assays were conducted using bacterial species that were identified in human breast tumors based on prior 16S rRNA sequencing data, including *Staphylococcus aureus*, *Staphylococcus epidermidis*, *Staphylococcus hominis*, *Streptococcus mitis*, *Streptococcus oralis*, *Cutibacterium acnes*, *Lactococcus lactis*, *Anaerococcus octavius*, *Bifidobacterium longum*, *Corynebacterium tuberculostearicum*, and *Pseudomonas aeruginosa*. Data from the 4T1 preclinical model of triple-negative breast cancer (TNBC) showed that, in most cases, intratumoral bacteria as modelled by intratumoral bacterial injections did not influence tumor growth (Fig. 5a). Only intratumoral injection of two *Staphylococcus* species, *S. aureus* or *S. hominis*, led to significant inhibition of tumor growth, with *S. aureus* displaying the most pronounced effect. This tumor-suppressive activity of *S. aureus* was corroborated in another TNBC model EO771, using PBS and *S. mitis* as negative controls (Fig. 5b).(35) Further investigation revealed that *S. aureus*-mediated inhibition of tumor growth was dependent on CD8^+^ T cells since this effect was nullified by antibody-mediated depletion of CD8^+^ cells but not CD4^+^ cells (Fig. 5c). Immunohistochemistry also showed that intratumoral *S. aureus* significantly increased GzmB-expressing cells in EO771 tumors, although it did not affect the abundance of CD8^+^ cells (Fig. 5d, e). Flow cytometry analysis confirmed *S. aureus* activity in stimulating GzmB expression in CD8^+^ cells in both EO771 and 4T1 models (Fig. 5f, g and supplementary Fig. 8). Beyond lymphocytes, intratumoral *S. aureus* modulated the innate immune cells in the tumor microenvironment. Without affecting dendritic cells, intratumoral *S. aureus* specifically reduced the tumor-associated macrophages, the ratio of M2-like (pro-tumoral) to M1-like (anti-tumoral) macrophages, and monocytic MDSCs (mMDSCs), but led to an increase in granulocytic myeloid-derived suppressor cells (gMDSCs) and polymorphonuclear leukocytes (PMNs) (supplementary Fig. 9a-e). Moreover, *S. aureus* appeared to attenuate the immunosuppressive functions of mMDSCs, gMDSCs, and PMNs by downregulating phosphorylated signal transducer and activator of transcription 3 (STAT3) (supplementary Fig. 9f, g).(36) Given significant correlation between intratumoral *Staphylococcus* and the cold metabolite NADH in human breast tumors (Fig. 3), we next investigated whether *S. aureus* functionally affects total NAD level. Indeed, intratumoral *S. aureus* but not *S. mitis* in mammary tumors led to a reduction in total NAD level within tumors (Fig. 5h). Overall, these results suggest that the presence of *S. aureus* in preclinical TNBC tumors depletes the cold metabolite NAD and activates CD8^+^ TILs, thereby enhancing anti-tumor immune responses, which is in concordant with positive correlations seen between levels of *Staphylococcus* and T cell activity in human breast tumors (Fig. 1c, 4e, 4f).

**Figure 5.**
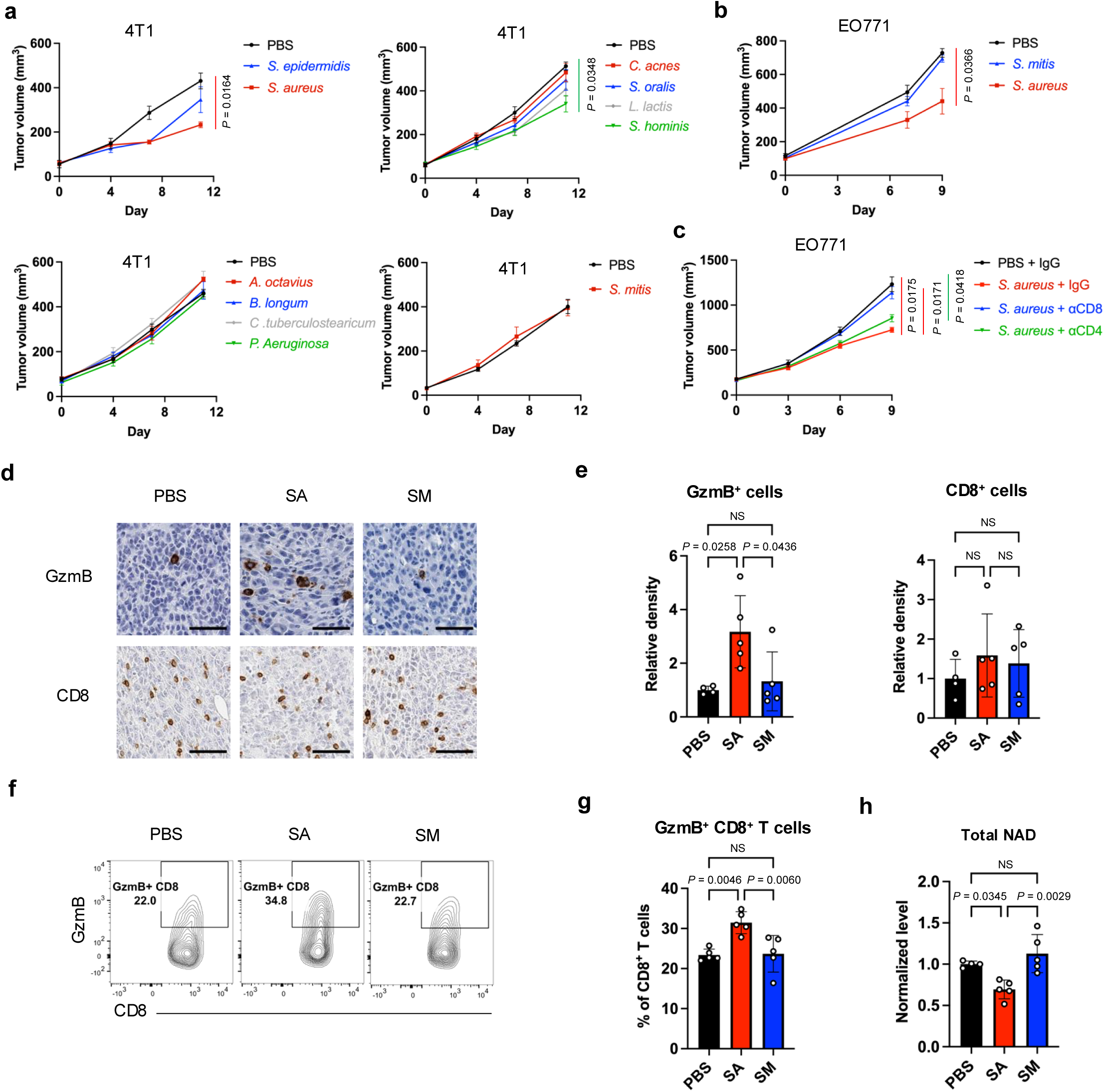
Intratumoral *S. aureus* modulates CD8^+^ TILs and total NAD level in TNBC models. a, **b** Growth of 4T1 (**a**) and EO771 (**b**) tumors after intratumoral injection of various bacterial species compared to PBS-treated control (n=3-5). **c** Growth of EO771 tumors after intratumoral injection of *S. aureus* with and without antibody-based depletion of CD8^+^ or CD4^+^ T cells (n=4). **d** Representative immunohistochemistry (IHC) images showing GzmB^+^ and CD8^+^ cells in EO771 tumors after intratumoral injection of *S. aureus* (SA) or *S. mitis* (SM), with PBS treatment as a control. Scale bars are shown in black at 50 μm. **e** Quantification of densities of GzmB^+^ and CD8^+^ cells based on IHC staining shown in **d**. **f** Representative contour plots of flow cytometric analysis showing the percentage of GzmB-expressing CD8+ T cells in EO771 tumors after intratumoral injection of *S. aureus* or *S. mitis*, with PBS treatment as a control. **g** Quantification of the percentage of GzmB+ cells among CD8^+^ T cells based on the flow cytometric plots shown in **f**. **h** Total levels of NAD+ and NADH in the EO771 tumors after intratumoral injection of *S. aureus* or *S. mitis*, with PBS treatment as a control. Two-way analysis of variance (ANOVA) with multiple comparisons (**a-c**). One-way ANOVA with multiple comparisons (**e, g, h**). NS, not significant.

## DISCUSSION

The role of the human breast tumor microbiome in breast cancer tumorigenesis, progression, and immune phenotypes remains understudied, partly due to challenges associated with the low microbial biomass in breast tissues and the difficulties of establishing causal relationships between the microbiome and cancer phenotypes. This study utilized two independent clinical cohorts, preclinical models, and multidimensional approaches to investigate the crosstalk between the breast tumor microbiome, anti-tumor immunity, and metabolome. We initially identified *Staphylococcus* as the only genus positively correlated with T cell activity in human breast tumors. Due to the small sample size, we did not stratify the analysis according to BC subtypes. To explore the potential functions of *Staphylococcus* across different BC subtypes, we subsequently analyzed RNA sequencing data from a larger cohort of 370 BC patients. This analysis revealed that positive association between *Staphylococcus* and T cell activity was exclusively present in TNBC. Accordingly, we further used preclinical TNBC models to demonstrate that intratumoral *S. aureus*, the predominant *Staphylococcus* species in human breast tumors, enhanced the expression of granzyme B in CD8^+^ TILs, thereby inhibiting tumor growth in a CD8^+^ cell-dependent manner. Notably, in human breast tissues, the abundance of *Staphylococcus* consistently showed a decreasing trend with advancing cancer stages across various breast cancer subtypes (Fig. 4b). Furthermore, the abundance of *Staphylococcus* was found to be lower in breast tumors compared to healthy breast tissues.(16) We therefore propose that *S. aureus* and potentially other *Staphylococcus* species may act as “sentinel bacteria” in breast tissues to protect against tumor development and progression by interacting with TILs. Similar tumor-suppressive activities of *S. aureus* have also been observed in glioblastoma, where the presence of intracranial *S. aureus* in several patients were associated with longer survival.(37) Future research is required to investigate whether the presence of *Staphylococcus* or *S. aureus* in TNBC influences responses to immunotherapy, as well as to characterize the functions of *Staphylococcus* in ER^+^/PR^+^ and other BC subtypes.

In addition to the tumor microbiome, deregulated cancer metabolism impairs TIL activities through the abnormal accumulation or depletion of immunomodulatory metabolites. This study systematically investigated the tumoral metabolites associated with CD8^+^ TILs using untargeted metabolome profiling and TIL analysis. Notably, our findings recaptured several metabolites previously identified to have immunomodulating functions in cancer. For example, among the hot metabolites, i.e., metabolites that are enriched in tumors with dense CD8^+^ TILs, linoleate and the microbiota-derived 3-formylindole (also called indole-3-aldehyde) were reported to stimulate cytotoxic T cell (CTL)-dependent anti-tumor immunity (Fig. 2d).(38, 39) Hot metabolites also include salicylate and 2-hydroxyhippurate (salicylurate), both of which are metabolic derivatives of aspirin (acetylsalicylic acid). Interestingly, aspirin intake was found to be associated with better responses to programmed death 1 (PD-1)/programmed death ligand 1(PD-L1)-based immunotherapy.(40, 41) As for cold metabolites, we showed that NAD^+^ and NADH are the most significant compounds that negatively correlate with both TIL abundance and T cell cytokines. Consistent with our findings, a previous study indicated that supplementation with NAD^+^ inhibited CD8^+^ TIL function.(42) The other cold metabolites, taurolithocholate 3-sulfate and glycolithocholate sulfate, are derivatives of secondary bile acids, whose functions in antagonizing CTL-mediated tumor cell killing were recently identified in colorectal cancer.(43) The cold metabolites also include sex hormone precursors DHEA-S and androstenediol (3beta,17beta) disulfate (2), which could be converted to testosterone and/or estrogen. Their desulfated forms DHEA and androstenediol (3beta,17beta) also retain weak estrogenic activity. Accordingly, estrogen signaling may be linked to the suppression of TIL function. While succinate was identified as a cold metabolite, other TCA cycle intermediates did not exhibit significant association with TILs, suggesting succinate-mediated TIL regulation might be independent of the alterations of TCA flux. In addition to these characterized metabolites, our study uncovered many TIL-associated compounds with unreported immunomodulatory functions, which are potential agents or targets for immunotherapy that require further validation.

In this study, we reported potential interactions between microbes and tumoral metabolites. For instance, the abundance of *Anaerococcus* was positively associated with an increased ratio of glutamate to glutamine, a potential indicator of glutaminolysis that is associated with breast cancer progression and TNBC (supplementary Fig. 4, 6).(34, 44) It is possible that enhanced tumoral glutaminolysis fosters an environment suitable for colonization by anaerobic *Anaerococcus* and other anaerobic bacteria, since the increased glutaminolysis could lead to hypoxia through oxidative phosphorylation-mediated oxygen consumption.(45) We also found that *Staphylococcus*-positive tumors exhibit reduced levels of GDP and the cold metabolite NADH, as well as increased abundance of hot metabolites including γ-glutamyltryptophan and γ-glutamylglutamate. It is likely that tumor-localized *Staphylococcus* directly consumes NADH and GDP to synthesize NADP(H) and alarmones (p)ppGpp, respectively, conferring survival advantage inside host cells similar to activity in other harsh conditions.(46, 47) Furthermore, we demonstrated in a preclinical BC model that intratumoral administration of *S. aureus*, but not *S. mitis*, led to a reduction of total NAD level in tumors, demonstrating that *Staphylococcus* plays a causal role in altering NAD and NADH.

While the precise means by which *S. aureus* activates CD8^+^ TILs in preclinical TNBC models remain unclear, we posit several potential mechanisms may be involved. *S. aureus* could modulate the CD8^+^ TIL-associated metabolites such as NAD and NADH and secrete various immunomodulatory agents, such as α-hemolysin, superantigens, and extracellular vesicles. In particular, α-hemolysin has been reported to stimulate CD8^+^ TILs and impede TNBC tumor growth.(30) Additionally, peptides from *S. aureus* can be presented on the surface of tumor cells in cancer patients, which has been demonstrated with activity to elicit CTLs.(48) Furthermore, *S. aureus* invasion could stimulate the STING signaling pathway by inducing DNA damage.(49) Future studies are needed to validate these and other crosstalk mechanisms.

Our study has several limitations that warrant consideration. First, the small sample size restricted the statistical power to detect more significant interactions between the tumor microbiome, TILs, and metabolites, which is a particular challenge in studying low-biomass tissues. Second, we were unable to culture patient tumor-derived *S. aureus* strains for functional studies due to the limited availability of tissue samples that had already been allocated for multiple analyses (microbiome, metabolome, TIL studies). Another constraint is that while our metabolome profiling showed high sensitivity that detected more than 900 intratumoral compounds, it did not provide insight into dynamic metabolic flux or specify the cell types contributing to metabolic alterations. Future studies should consider incorporating isotope-labeling techniques and integrating further omics analyses, including single-cell RNA sequencing, high-plex RNAscope-FISH, high-plex immunofluorescence imaging, spatial transcriptomics, spatial metabolomics, and tumor culturomics to address these limitations.

### CONCLUSIONS

This study identifies microbial taxa and metabolites within human breast tumors that are associated with CD8^+^ TILs. We also explore crosstalk between the tumor microbiome and metabolome, and demonstrate that the presence of *Staphylococcus* within tumors stimulates CD8^+^ TIL-dependent anti-tumor immunity in TNBC and modulates T cell-associated metabolites. These findings illuminate the potential significant roles of the low-biomass tumor microbiome in anti-tumor immunity and cancer metabolism, and lay the groundwork for further studies assessing whether these CD8^+^ TIL-associated microbial taxa and metabolites could serve as biomarkers or therapeutic agents for immunotherapy, increasing its efficacy in TNBC or expanding its usage in other BC subtypes.

## LIST OF ABBREVIATIONS

BC: Breast cancer
CCA: Canonical correlation analysis
CFU: Colony forming units
CTL: Cytotoxic T cell
DHEA-S: Dehydroepiandrosterone sulfate
ER: Estrogen receptor
ESI: Electrospray ionization
FISH: Fluorescence *in situ* hybridization
gMDSC: Granulocytic myeloid-derived suppressor cell
HER2: Human epidermal growth factor receptor 2
ICI: Immune checkpoint inhibitor
mMDSC: Monocytic myeloid-derived suppressor cell
NAAG: N-acetyl-aspartyl-glutamate
NAD: Nicotinamide adenine dinucleotide
NADH: Nicotinamide adenine dinucleotide hydrogen
PBS: Dulbecco’s phosphate-buffered saline
PD-1: Programmed death 1
PD-L1: Programmed death ligand 1
PMN: Polymorphonuclear leukocyte
PR: Progesterone receptor
STAT3: Signal transducer and activator of transcription 3
TIL: Tumor-infiltrating lymphocyte
TNBC: Triple-negative breast cancer
UPLC-MS/MS: Ultrahigh performance liquid chromatography-tandem mass spectroscopy

## DECLARATIONS

### Ethics approval and consent to participate

Human tissue and data for this study were collected under Cleveland Clinic Institutional Review Board-approved protocols IRB #14-774 and 17-791, in which written informed consent was obtained from adult research participants in alignment with the Helsinki Declaration. All protocols of animal experiments were approved by the Institutional Animal Care and Use Committee (IACUC) of Cleveland Clinic (protocol number: 00002429). All experimental methods were carried out in accordance with the approved protocol and the guidelines and regulations of IACUC. Animal experiments were performed and reported in accordance with Animal Research: Reporting of In Vivo Experiments (ARRIVE) guidelines. The Ohio State University Institutional Review Board (IRB) approved data access in an Honest Broker protocol (2015H0185) and Total Cancer Care protocol (2013H0199) in coordination with Aster Insights.

### Consent for publication

Not applicable.

### Availability of data and materials

The data generated or analyzed during this study are included in this published article and its supplementary information files. Custom code scripts are available in the code repository https://github.com/spakowiczlab/staph-brca.

## Contributions

C.L. and C.E. conceptualized the study. S.R.G. and Z.A. obtained surgical samples of Cohort A. C.L., D.G., R.L., and M.W. carried out the experiments. C.L., K.C., Y.N., L.L., R.H., M.W., and N.S. analyzed the data and/or performed statistical analyses. C.L., C.E., Y.N., D.S., K.C., D.G., and S.R.J. interpreted the analyses. A.P.S. validated the results of T cell identification analyzed using the CaloPix software. C.L. drafted the manuscript. A.T., D.S., D.G., Y.N., S.R.G., S.R.J., and K.C. critically revised the manuscript. C.E., Y.N., and D.S. oversaw the project. All authors except the late C.E. read and approved the final manuscript.

## Acknowledgments

We appreciate the patients who generously donated their samples to this project. We thank Tammy Sadler and Qi Yu (Eng lab) for assisting with this study. We thank Diana Moorman and Eric Schultz of the Flow Cytometry Core (Cleveland Clinic Lerner Research Institute, CC-LRI), Andrelie Branicky and Zalavadia Ajay of the Imaging Core (CC-LRI), Microbial Culturing and Engineering Core (CC-LRI), and Saima Ben Hadj and Tribun Healthcare for their technical assistance.

