## Supplementary Figures and Tables for "Breast tumor microbiome regulates anti-tumor immunity and T cell-associated metabolites"

**Supplementary figures and legends 1 to 10**

**Supplementary tables 1 and 2**

### Supplementary figure 1

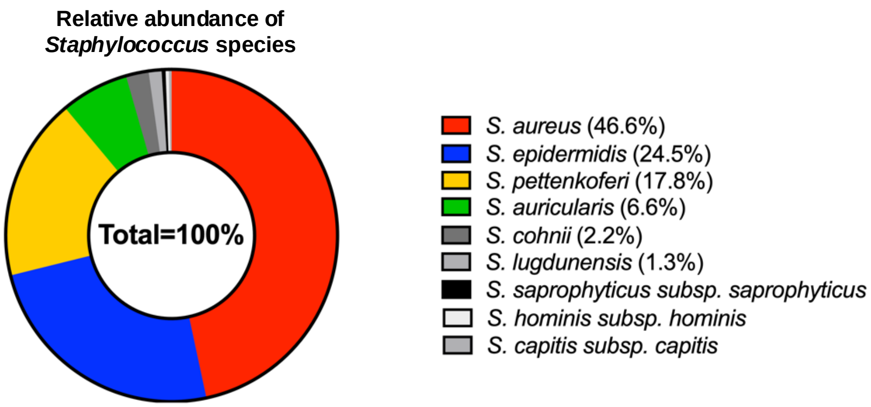


**Supplementary figure 1 | The relative abundance of *Staphylococcus* species in human breast tumors.** The percentage of *Staphylococcus* species in human breast tumors based on the reads identified in 16S rRNA gene sequencing.

### Supplementary figure 2

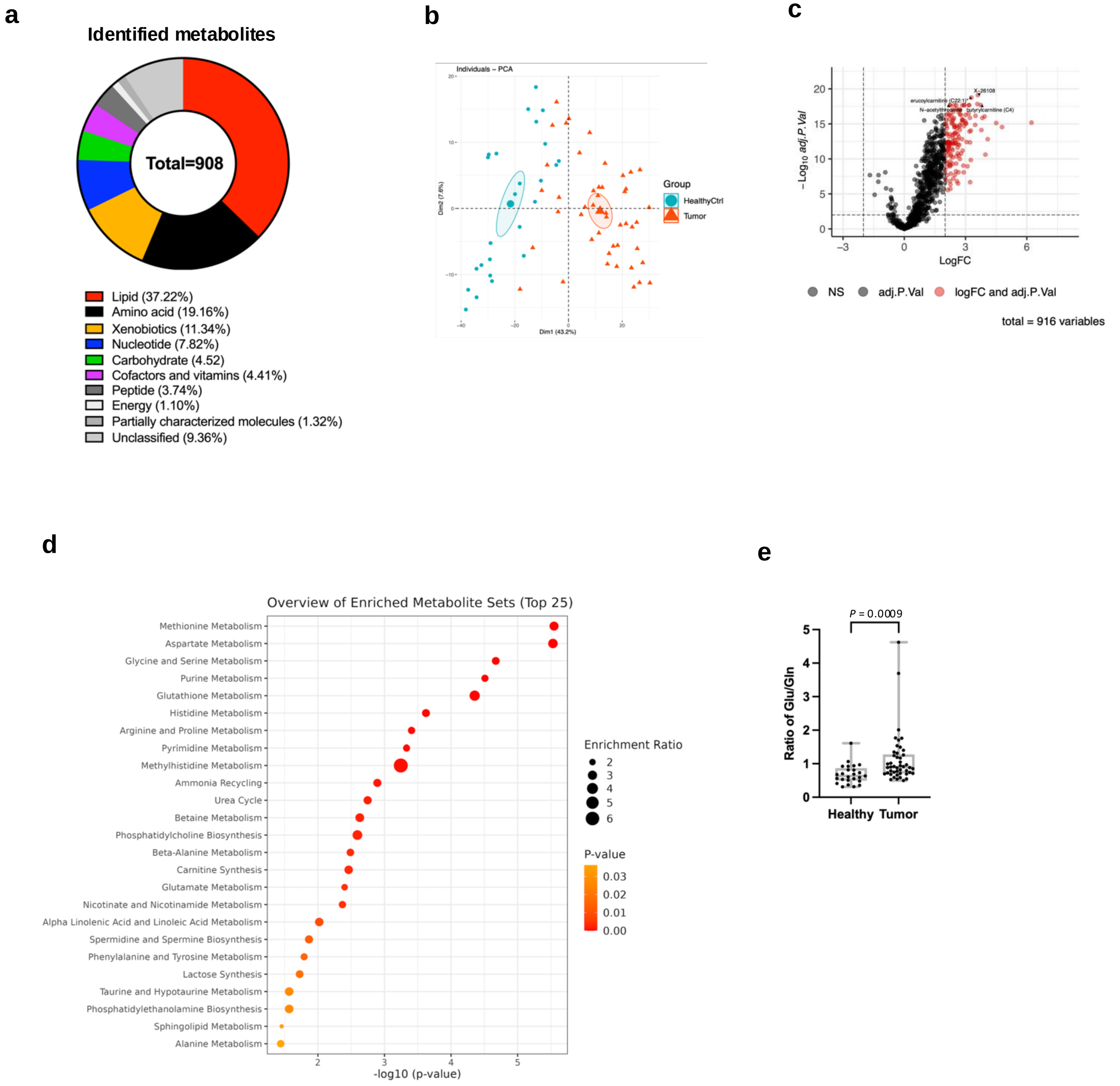


**Supplementary figure 2 | Metabolic differences between breast tumors and non-malignant breast tissues. a** Composition of the metabolites identified by untargeted metabolomics in human breast tumors and healthy breast tissues. **b** Principal component analysis (PCA) of metabolites in human breast tumors (n=46, in orange) and healthy breast tissues (n=25, in blue). **c** The volcano plot showing the differentially abundant metabolites between breast tumors and healthy breast tissues. Metabolites with a log2 fold change > 2 and –Log10 adjusted p-value > 1.3 are highlighted in red. **d** Top metabolic pathways that are significantly altered in breast tumors compared to healthy breast tissues. **e** Distinct ratios of glutamate (Glu) to glutamine (Gln) between breast tumors and healthy breast tissues.

### Supplementary figure 3

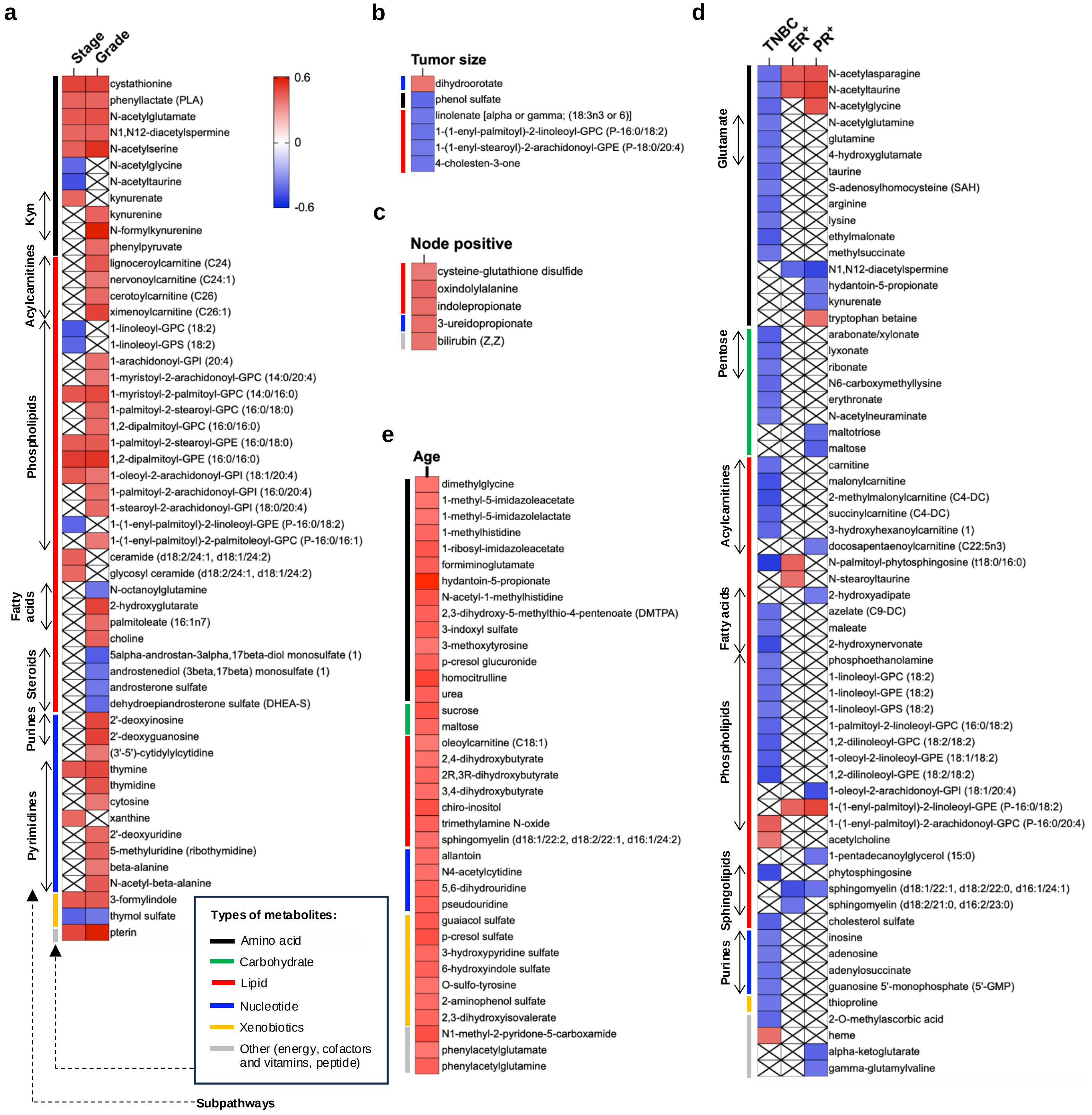


**Supplementary figure 3 | Associations between metabolites and clinicopathological features of breast cancer. a-e** Heatmaps showing the metabolites significantly associated with cancer stage or histological grade (**a**), tumor size (**b**), lymph node-positive status (**c**), histologic subtypes (**d**), and patient age (**e**) with the color gradient representing correlation. Types and subpathways of metabolites are indicated on the left side of each heatmap.

### Supplementary figure 4

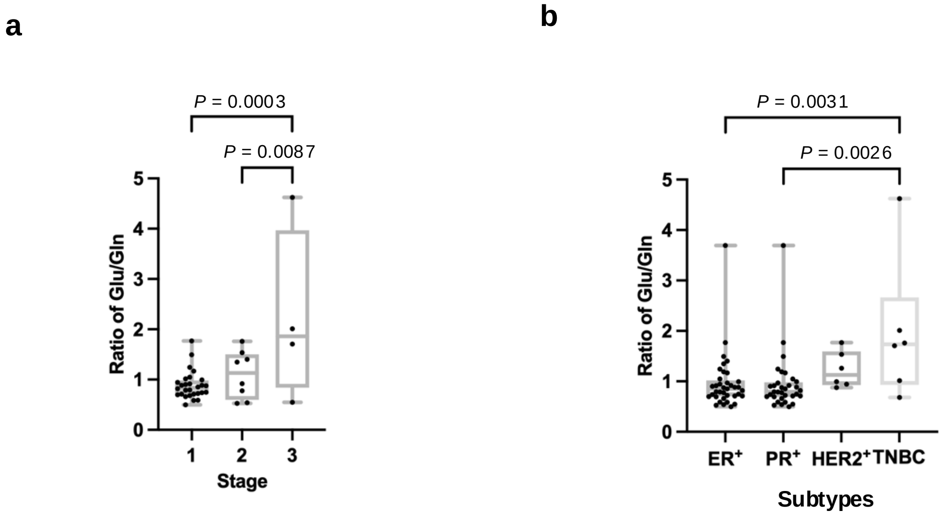


**Supplementary figure 4 | The ratio of glutamate to glutamine differs between BC stages and subtypes. a, b** Ratio of glutamate (Glu) to glutamine (Gln) in breast tumors across different stages (**a**) and subtypes (**b**). One-way analysis of variance (ANOVA) with multiple comparisons. Only the significant differences are indicated with p-values.

### Supplementary figure 5

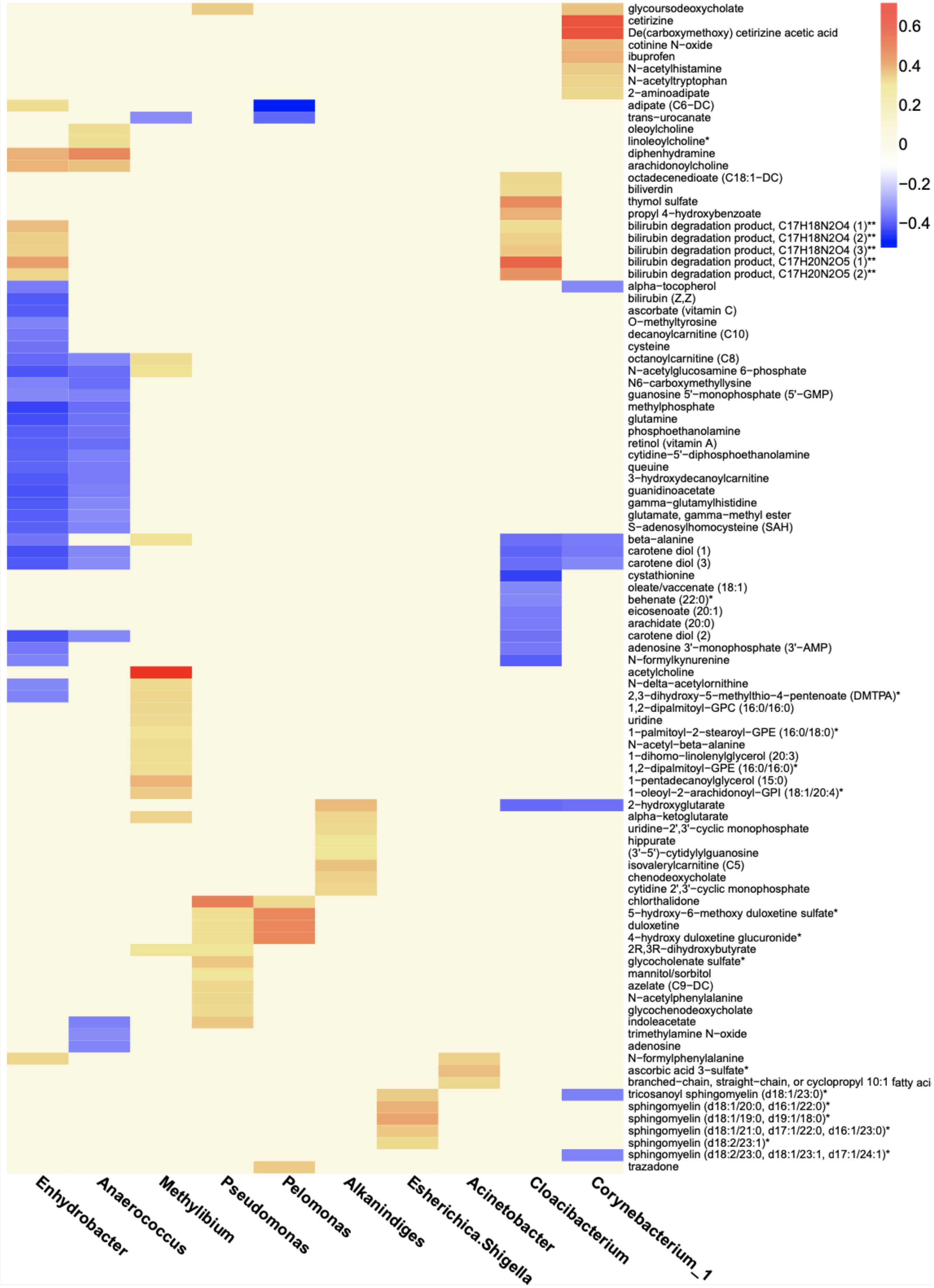

**Supplementary figure 5 | Correlations between tumoral bacteria and metabolites.** Heatmap displaying significant correlations between the bacterial genera and metabolites within human breast tumors identified by sparse canonical correlation analysis (CCA). The color gradient representing the correlation.

### Supplementary figure 6

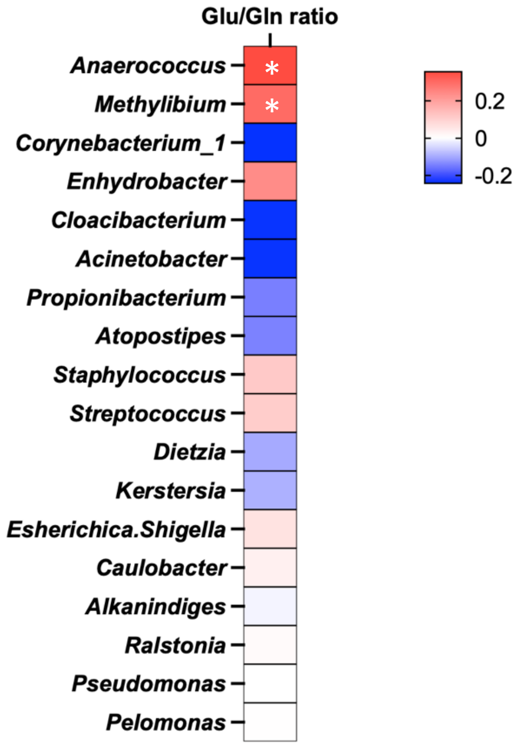


*

**Supplementary figure 6 | Correlations between tumoral bacteria and the ratio of glutamate to glutamine.** Heatmap illustrating the correlations between bacterial genera and the ratio of glutamate (Glu) to glutamine (Gln). Spearman's rank correlation. *$P < 0.05$.

### Supplementary figure 7

**a**

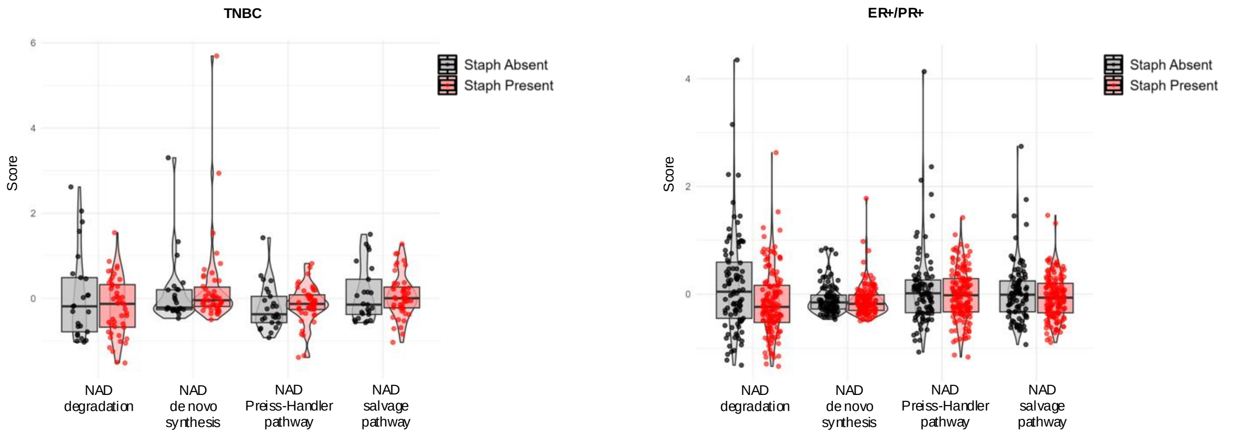


**b**

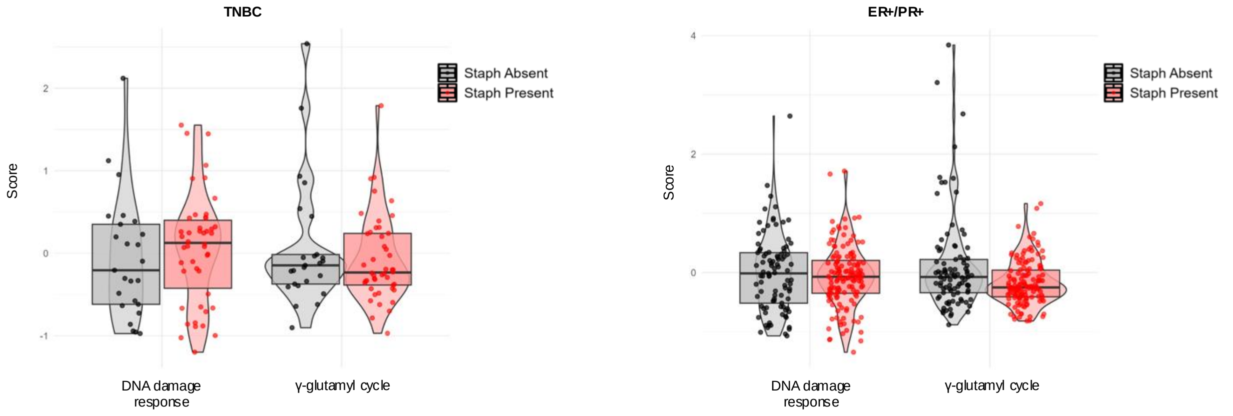


**Supplementary figure 7 | Associations between *Staphylococcus* and metabolic pathways in breast cancer. a, b** Comparisons of the activity of NAD-related pathways (**a**), DNA damage response (**b**), and γ-glutamyl cycle pathways (**b**) in TNBC (left panel) and ER+/PR+ subtype (right panel) with and without *Staphylococcus* (Staph, denoted in red and black dots, respectively). Z-score transformed signature scores were compared by t-test. All p-values were > 0.05.

### Supplementary figure 8

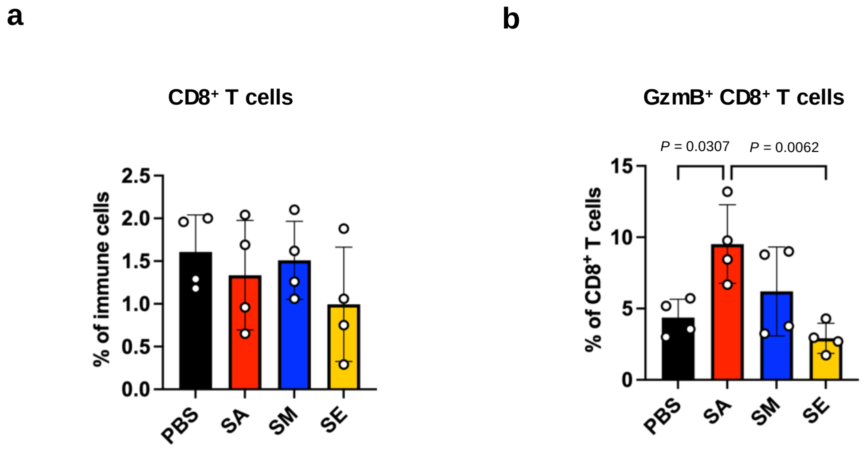


**Supplementary figure 8 | Tumoral colonization by *S. aureus* activates CD8+ T cells in 4T1 tumors. a, b** 4T1 tumors were colonized by *S. aureus* (SA), *S. mitis* (SM), or *S. epidermidis* (SE), with PBS treatment as a control. Flow cytometry was performed to determine the percentage of $CD8^+$ T cells among immune cells (**a**) and $GzmB^+$ $CD8^+$ T cells among $CD8^+$ T cells (**b**). Only significant differences are indicated with *p*-values. One-way analysis of variance (ANOVA) with multiple comparisons.

### Supplementary figure 9

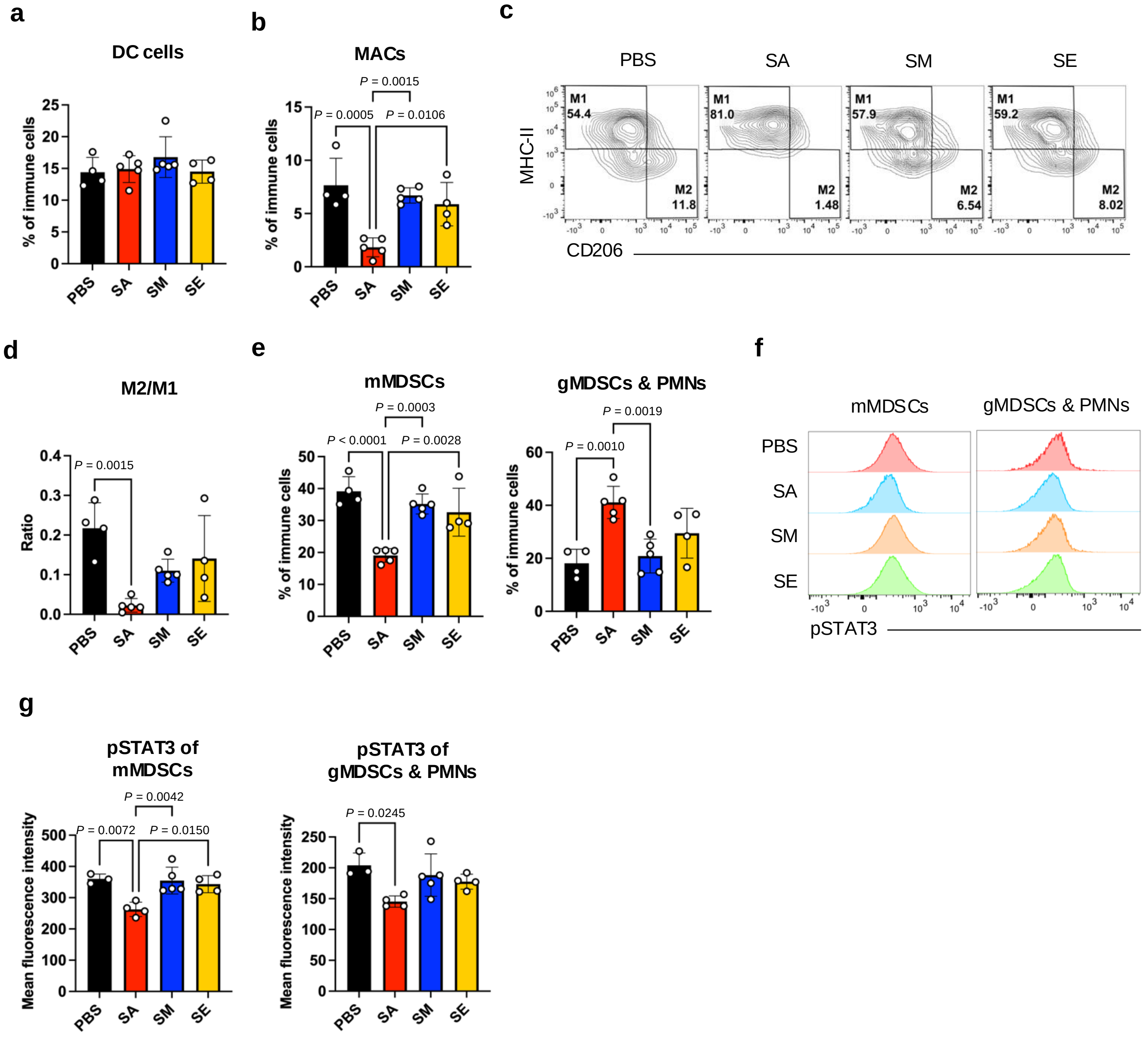


**Supplementary figure 9 | Tumoral colonization by *S. aureus* affects innate immune cells in EO771 tumors. a, b, d, e, g** Flow cytometry analysis of innate immune cells in the EO771 tumors colonized by *S. aureus* (SA), *S. mitis* (SM), or *S. epidermidis* (SE), with PBS treatment as a control. The analyses including the percentage of dendritic cells (DCs) among immune cells (**a**), the percentage of macrophages (MASc) among immune cells (**b**), the ratio of M2-like to M1-like MACs (**d**), the percentage of monocytic myeloid-derived suppressor cells (mMDSCs) and granulocytic MDSC (gMDSCs)/polymorphonuclear neutrophils (PMNs) among immune cells (**e**), and the mean fluorescence intensity of phosphorylated STAT3 in mMDSCs and gMDSCs/PMNs (**g**). **c**, Representative contour plots showing the gating of M2-like (MHC-II$^{low}$, CD206$^{+}$) and M1-like (MHC-II$^{high}$, CD206$^{-}$) MACs in the tumors. **f** Representative histograms showing the levels of phosphorylated STAT3 in mMDSCs and gMDSCs/PMNs. One-way analysis of variance (ANOVA) with multiple comparisons (**a**, **b**, **d**, **e**, **g**). Only the significant differences were denoted with *p*-values.

### Supplementary figure 10

**a**

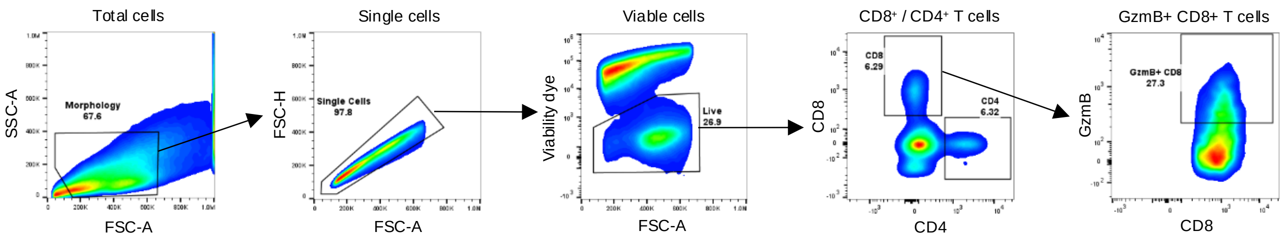


**b**

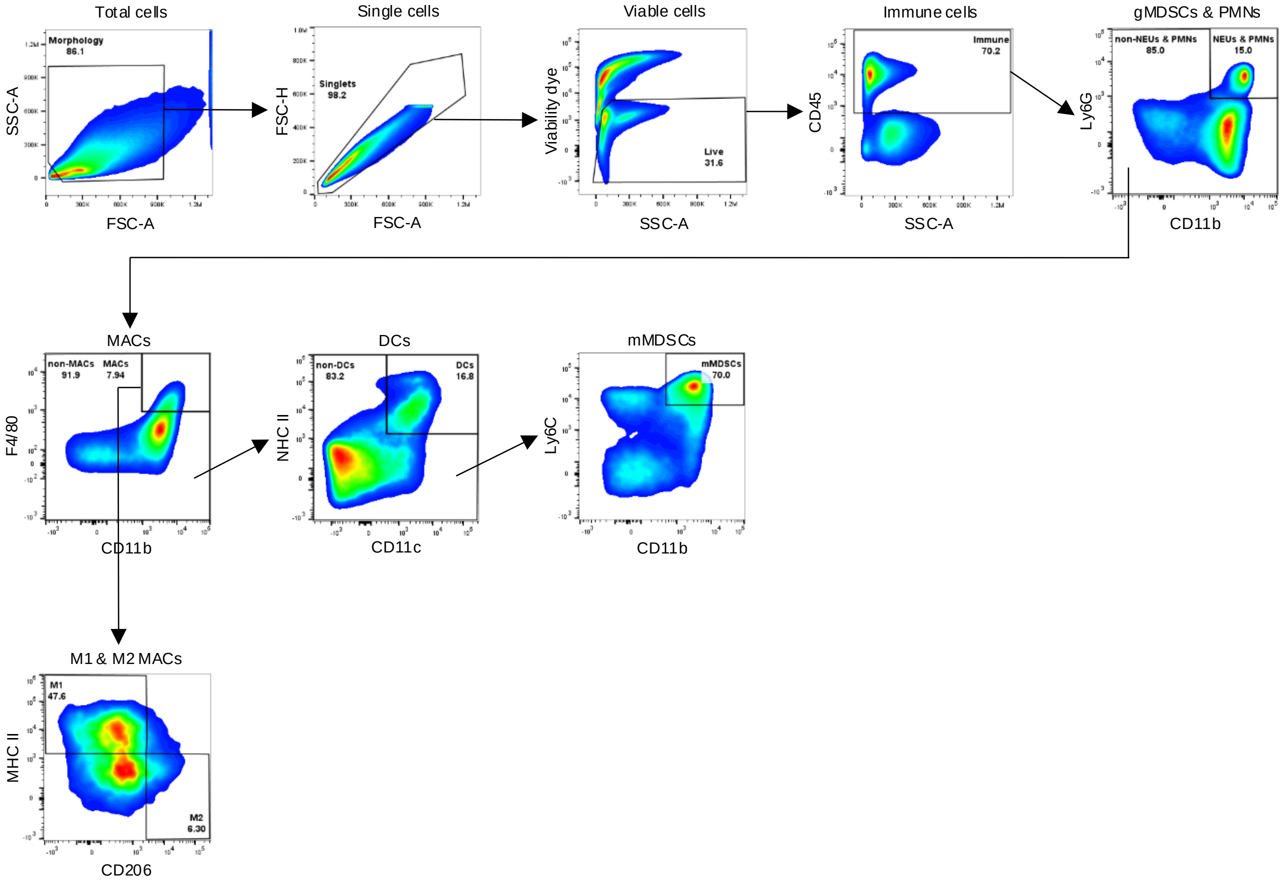


**Supplementary figure 10 | Gating strategy for flow cytometry used throughout the manuscript. a, b** Gating strategy for GzmB⁺ CD8⁺ T cells (**a**) and granulocytic myeloid-derived suppressor cells (gMDSCs) and polymorphonuclear leukocytes (PMNs), macrophages (MACs), M1- and M2-like MACs, dendritic cells (DCs), and monocytic MDSCs (mMDSCs) (**b**).

**Supplementary Table 1 | The top 25 metabolites significantly enriched in breast tumors compared to healthy breast tissues.**

| Metabolites enriched in control breast tissues | Log2 fold change (tumor/control) | Adjusted p-value |
|---|---|---|
| erucoylcarnitine (C22:1)* | 3.28 | 1.75E-19 |
| arachidoylcarnitine (C20)* | 3.58 | 1.45E-18 |
| nervonoylcarnitine (C24:1)* | 3.16 | 2.11E-18 |
| docosadienoylcarnitine (C22:2)* | 3.18 | 2.11E-18 |
| butyrylcarnitine (C4) | 3.74 | 2.11E-18 |
| behenoylcarnitine (C22)* | 3.15 | 3.40E-17 |
| glycerophosphoethanolamine | 3.01 | 5.36E-17 |
| glutamate, gamma-methyl ester | 3.49 | 6.99E-17 |
| 5-methylthioadenosine (MTA) | 3.93 | 8.58E-17 |
| eicosenoylcarnitine (C20:1)* | 3.22 | 1.35E-16 |
| N-acetylaspartate (NAA) | 3.82 | 1.45E-16 |
| ascorbate (vitamin C) | 6.23 | 6.55E-16 |
| phytosphingosine | 4.50 | 7.50E-16 |
| N-acetylputrescine | 3.00 | 8.86E-16 |
| N-palmitoyl-phytosphingosine (t18:0/16:0) | 3.47 | 1.39E-15 |
| alpha-tocopherol | 4.79 | 2.80E-15 |
| cytidine 5'-diphosphocholine | 3.78 | 7.57E-15 |
| quinolinate | 3.26 | 9.18E-15 |
| ethylmalonate | 3.19 | 1.23E-14 |
| UDP-N-acetylglucosamine/galactosamine | 4.08 | 1.52E-13 |
| N-acetyl-aspartyl-glutamate (NAAG) | 3.05 | 3.85E-13 |
| cystathionine | 3.97 | 2.43E-11 |
| guanosine 5'-monophosphate (5'-GMP) | 3.26 | 1.44E-10 |
| uridine 5'-monophosphate (UMP) | 3.09 | 1.15E-09 |
| glutathione, reduced (GSH) | 3.23 | 2.10E-07 |

Mann-Whitney U test

**Supplementary Table 2 | The top 25 metabolites significantly enriched in healthy breast tissues compared to breast tumors.**

| Metabolites enriched in control breast tissues | Log2 fold change (tumor/control) | Adjusted p-value |
|---|---|---|
| sphingomyelin (d18:1/20:1, d18:2/20:0)* | -0.93 | 8.33E-09 |
| trans-urocanate | -1.68 | 2.07E-08 |
| linolenate [alpha or gamma; (18:3n3 or 6)] | -0.89 | 1.64E-06 |
| sphingomyelin (d18:1/18:1, d18:2/18:0) | -0.65 | 2.71E-06 |
| sphingomyelin (d18:2/24:2)* | -0.65 | 3.07E-05 |
| 3-hydroxy-2-methylpyridine sulfate | -0.63 | 5.93E-05 |
| caprylate (8:0) | -0.60 | 0.00011136 |
| bilirubin degradation product, C17H20N2O5 (2)** | -0.70 | 0.00030644 |
| octadecadienedioate (C18:2-DC)* | -0.79 | 0.00051615 |
| branched-chain, straight-chain, or cyclopropyl 10:1 fatty acid (1)* | -0.61 | 0.00087754 |
| triethanolamine | -0.63 | 0.00105426 |
| o-cresol sulfate | -0.65 | 0.00128545 |
| 13-HODE + 9-HODE | -0.47 | 0.00345925 |
| 1-(1-enyl-palmitoyl)-2-linoleoyl-GPC (P-16:0/18:2)* | -0.58 | 0.00820577 |
| androstenediol (3beta,17beta) disulfate (1) | -0.79 | 0.00902972 |
| 3-amino-2-piperidone | -0.64 | 0.00986265 |
| octadecenedioate (C18:1-DC) | -0.54 | 0.01034876 |
| sphingomyelin (d18:2/18:1)* | -0.52 | 0.01066854 |
| 4-vinylphenol sulfate | -0.79 | 0.0109589 |
| N-behenoyl-sphingadienine (d18:2/22:0)* | -0.39 | 0.01771484 |
| 12,13-DiHOME | -0.54 | 0.0190684 |
| 5alpha-androstan-3beta,17beta-diol disulfate | -0.66 | 0.02234858 |
| fructosyllysine | -0.76 | 0.02345034 |
| chenodeoxycholate | -0.56 | 0.03225341 |
| 2-aminophenol sulfate | -0.48 | 0.04465686 |

Mann-Whitney U test
